# Fetal Liver-like Organoids Recapitulate Blood-Liver Niche Development and Multipotent Hematopoiesis from Human Pluripotent Stem Cells

**DOI:** 10.1101/2024.10.11.617794

**Authors:** Milad Rezvani, Kyle Lewis, Susanna Quach, Kentaro Iwasawa, Julian Weihs, Hasan Al Reza, Yuqi Cai, Masaki Kimura, RanRan Zhang, Yuka Milton, Praneet Chaturvedi, Konrad Thorner, Ramesh C. Nayak, Jorge Munera, Phillip Kramer, Brian R. Davis, Appakalai N. Balamurugan, Yeni Ait Ahmed, Marcel Finke, Rose Yinghan Behncke, Adrien Guillot, René Hägerling, Julia K. Polansky, Philip Bufler, Jose A Cancelas, James M. Wells, Momoko Yoshimoto, Takanori Takebe

**Author notes:** Contributed equally (Milad Rezvani, Kyle Lewis, Susanna Quach).

## Abstract

The fetal liver is a hematopoietic organ, hosting a diverse and evolving progenitor population. While human liver organoids derived from pluripotent stem cells (PSCs) mimic aspects of embryonic and fetal development, they typically lack the complex hematopoietic niche and the interaction between hepatic and hematopoietic development. We describe the generation of human Fetal Liver-like Organoids (FLOs), that model human hepato-hematopoietic interactions previously characterized in mouse models. Developing FLOs first integrate a yolk sac-like hemogenic endothelium into hepatic endoderm and mesoderm specification. As the hepatic and hematopoietic lineages differentiate, the FLO culture model establishes an autonomous niche capable of driving subsequent progenitor differentiation without exogenous factors. Consistent with yolk sac-derived waves, hematopoietic progenitor cells (HPCs) within FLOs exhibit multipotency with a preference for myeloid lineage commitment, while retaining fetal B and T cell differentiation potential. We reconstruct in FLOs the embryonic monocyte-to-macrophage and granulocyte immune trajectories within the FLO microenvironment and assess their functional responses in the liver niche. *In vivo*, FLOs demonstrate a liver engraftment bias of hematopoietic cells, recapitulating a key phenomenon of human hematopoietic ontogeny. Our findings highlight the intrinsic capacity of liver organoids to support hematopoietic development, establishing FLOs as a platform for modeling and manipulating human blood-liver niche interactions during critical stages of development and disease.

## INTRODUCTION

The human fetal liver is a hematopoietic niche constituted by hepatobiliary cells and a complex non-parenchymal population. This niche is filled with a heterogeneous hematopoietic cell population that evolves and shifts its differentiation output throughout development (Popescu et al., 2019; Vanuytsel et al., 2022; Wesley et al., 2022). Hematopoietic stem and progenitor cells (HSPCs) from various intra- and extraembryonic sites engraft in the liver, where they undergo maturation or immune cell differentiation (Calvanese et al., 2022; Canu & Ruhrberg, 2021; Goh et al., 2023). After the initial primitive hematopoiesis from the yolk sac, a second wave of HSPCs from the yolk sac has been reported to populate the developing mouse liver with erythromyeloid and lymphomyeloid lineages (Canu & Ruhrberg, 2021; Kobayashi et al., 2023; Yokomizo et al., 2022). In human development, the transition from the yolk sac to liver hematopoiesis can be tracked by the hemoglobin (HB) subtypes, shifting from HBZ/HBE- to HBG1-predominant because the liver suppresses embryonic erythropoiesis (Goh et al., 2023). Following Carnegie Stage 14 (CS14) in human development, particularly after establishing the aorta–gonad–mesonephros (AGM) region, early HSPCs with an erythromyeloid bias emerge, albeit retaining some lymphoid potential. Subsequently, late yolk sac-derived HSPCs with a lymphoid and megakaryocyte bias become evident (Goh et al., 2023). Another study lends well to said findings, demonstrating a dynamically shifting hematopoietic heterogeneity in the liver from granulocyte–monocyte progenitors (GMPs) and lymphomyeloid progenitors at CS12 to more lymphoid-biased progenitors at CS15 (Bian et al., 2020). These findings indicate that human liver hematopoiesis transitions from myeloid to lymphoid bias. In mice, multipotent HSPCs of the pro-definitive wave of hematopoiesis are thought to only transiently reside in the bone marrow (Canu & Ruhrberg, 2021).

Progress has been made in modeling extraembryonic hematopoiesis *in vitro*, e.g., yolk sac hematopoiesis (Atkins et al., 2022). A recent study demonstrated the breakthrough of AGM-like differentiated human induced pluripotent stem cells (hiPSCs) that give rise to induced HSPCs engrafting the bone marrow of immune-deficient mice (Ng et al., 2024). Further, a bone marrow-like organoid from hiPSCs has been shown, giving rise to multipotent HSPCs with bone marrow engraftment capacity (Frenz-Wiessner et al., 2024). Different protocols exist to generate human embryonic or fetal hepatocytes or liver organoids (Hu et al., 2018; Shinozawa et al., 2021; Takebe et al., 2015; Wesley et al., 2022; Zhu et al., 2014). However, existing liver models lack the integration of a complex and dynamically changing hematopoietic system and niche-constituting cells, including fetal hepatocytes and mesenchymal cells, which provide the essential signaling crosstalk across progenitors at the fetal liver stage (Soares-da-Silva et al., 2020). Such an integrated model of hematopoietic and hepatic lineages – especially if it recapitulated hematopoietic cells seeding the fetal liver – could help understand early blood and co-liver development in humans and the clinical significance for hematopoietic or certain newborn liver disorders. Further, a better understanding of human hematopoiesis independent of definitive hematopoietic stem cells (HSCs) may be of translational value because in mice, HSC-independent hematopoietic progenitors have been shown to contribute significantly to adult mouse blood composition (Kobayashi et al., 2023; Patel et al., 2022)

Developing a human hepato-hematopoetic system is a complex task due to the contribution of multiple lineages and dynamic tissue crosstalk within the hematopoietic niche. Single-cell genomic datasets indicate the diversity and critical role of mesenchyme in endoderm organogenesis (Han et al., 2020). Studies have evolved multiple protocols to generate and recapitulate more complex epithelial-mesenchymal interactions. For example, progenitors from different germ layers can be differentiated separately and coaxed to develop multilineage tissues in the gut and liver organoids (Workman et al., 2017) (Múnera et al., 2023). Studies detected macrophages and stromal cell types in human liver organoids made from PSCs (Guan et al., 2021; Ouchi et al., 2019).

Herein, we demonstrate the generation of human Fetal Liver-like Organoids (FLOs), enabling the functional analysis of complex hepato-hematopoietic interactions described in mice or human fetal liver atlas. We show the essential interactive co-development of endoderm and mesoderm trajectories within FLOs. This results in the autonomous establishment of a niche capable of supporting blood and liver differentiation without reliance on exogenous differentiation factors. The hematopoietic progenitor cell (HPC) population within FLOs is heterogeneous and exhibits multipotency, favoring myeloid lineage commitment while retaining fetal B and T cell potential. Following the transplantation of FLOs into mice, myeloid cell engraftment in the liver was observed, mirroring *in vivo* developmental liver population by yolk sac-derived monocytes and macrophages. Moreover, we elucidate the fetal liver monocyte trajectories to macrophage and granulocyte immunity within the framework of a liver niche and determine their functional response.

## RESULTS

### Generating Fetal Liver-like Organoids (FLOs) with Integrated Hematopoiesis

We aimed to establish an organoid model integrating a hemogenic lineage into a three-dimensional fetal liver context (Fig. 1A). We co-induced endoderm (mean efficiency = 72.15%, SD ± 18.8) and yolk sac mesoderm (mean efficiency = 3.36, SD ± 1.97) using Activin A (ActA), BMP4 and FBS (Fig. 1B, C). We generated FLOs from n=10 different hPSC lines at three independent laboratories (Supplemental Table 1). To mimic signaling from cardiac mesoderm and septum transversum mesenchyme on hepatic differentiation, we added BMP4 and FGF2. These factors promoted the differentiation of the epithelial compartment and the development of CD34+ cells (Fig. 1D). From day 10, we let FLOs self-assemble in clusters without exogenous cytokines. In the analogy of developing human livers populated with macroscopically visible red blood cells from CS14 (post-conception week, pcw 5), erythroid islands colonize previously pale FLOs from day 14. These appeared as multiple colonies or bound within the boundaries of evolving reticular endothelial structures (Fig. 1E). Whole FLO light sheet microscopy demonstrates an ECAD+ epithelial architecture pervaded with a CD34+CD31+ hemogenic network and an apparent subset of CD34+ CD31-putative HPCs (Supplementary Video 2, Fig. 1F, G). Gene expression over time revealed that erythropoietin (*EPO*) trajectory followed albumin (*ALB*) along with alpha-fetoprotein (*AFP*) from day 6 (Figure 1H, Supplementary Table 2). Live cell microscopy revealed that clusters of progenitor-like cells acquired a red color near epithelial structures (Supplemental Video 1). RUNX1 is a marker for hematopoietic cells, such as HSPCs, that originate from hemogenic endothelium (Choi et al., 2011; North et al., 2002). RUNX1-expressing HSPCs are commonly found in the human fetal liver from pcw 4.5 (Calvanese et al., 2022). In FLOs, a surge in *RUNX1* expression from day 14 preceded erythroid organoid colonization (Figure 1H). We developed human embryonic stem cells (ESCs) with a *RUNX1+23* enhancer-reporter (Fig. S1A)(Bee et al., 2009). Reporter activity was detected before the onset of erythropoiesis from day 12 and continued in newly emerged red blood cell colonies on day 20 (Fig. 1I). When FACS-isolated for GYPA+, erythroid cells demonstrated nucleated erythroid morphology in cytological analysis (Fig. S1B). By multiplexed immunofluorescence (mIF), we identified CD34+ cells, CD31+ endothelial cells, IBA1+ macrophages, and HepPar1+ hepatocyte-like cells (Fig. 1J). HepPar1+ cells were present for over 40 days, while there was an inverse relationship between the increase in macrophages after day 24 and the gradual decrease of CD34+ cells (Fig. S1C). Immunohistochemistry (IHC) of FLOs and primary human fetal livers demonstrated that FLOs recapitulate AFP+/Albumin+ fetal hepatocytes. However, parenchymal densities in FLOs are lower than in primary fetal livers (Fig. S1D). Thus, FLOs promote hepatocellular and non-parenchymal cell differentiation during the initial stage without extrinsic factors after lineage specification. Moreover, they integrate a yolk sac mesoderm-derived hematopoietic lineage, progressively establishing hematopoietic activity with a transition from erythroid output to myeloid from certain types of progenitors.

**Figure 1.**
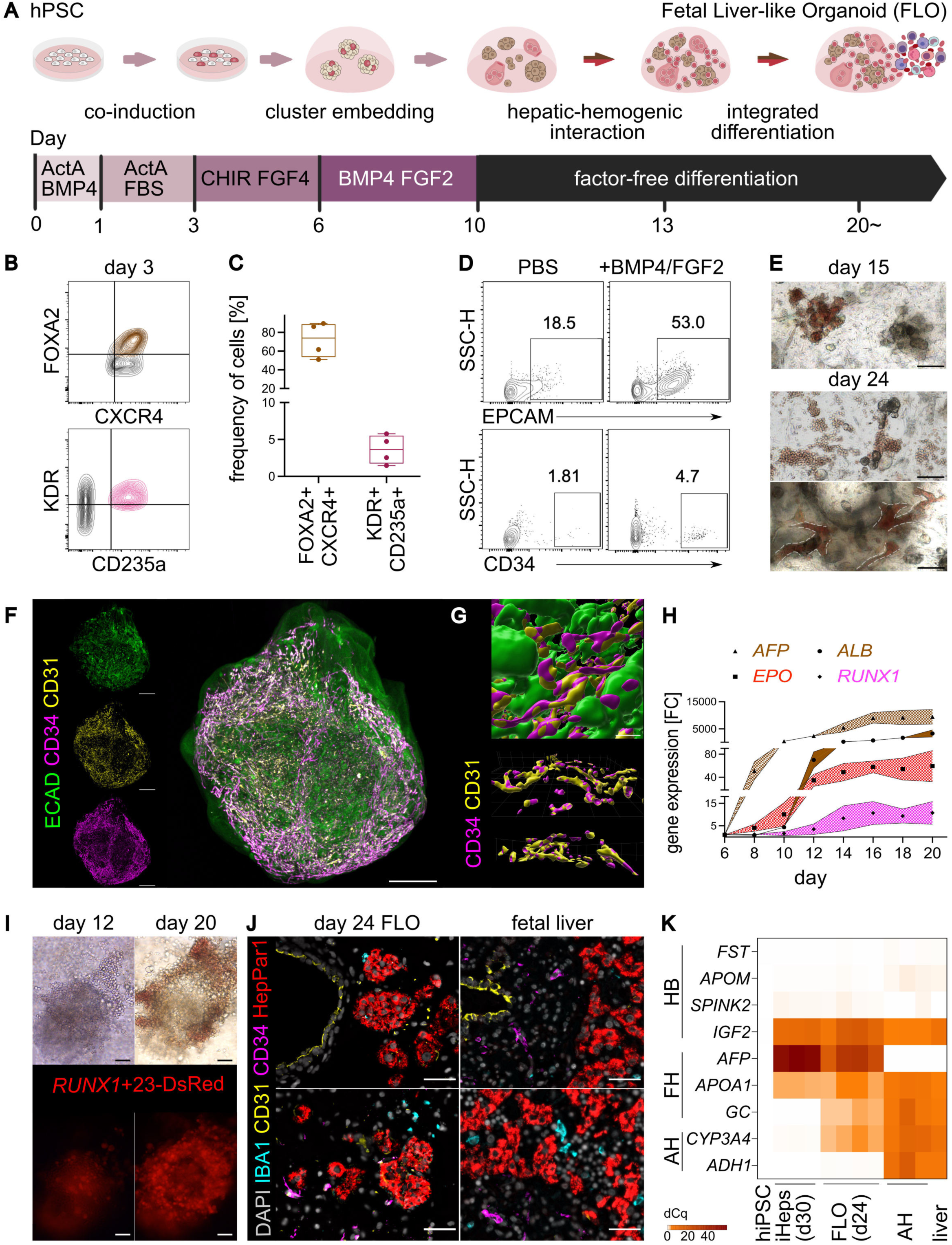
Generating Fetal Liver-like Organoids with Integrated Hematopoiesis. **A**: Schematic of generating fetal liver-like organoids (FLOs) from human pluripotent stem cells (hPSCs) through an initiation phase until day 10, followed by factor-free differentiation. ActA = Activin A. **B**: Representative flow cytometry of FOXA2+ CXCR4+ endoderm and KDR+ CD235a+ hemogenic mesoderm on day 3 of FLO differentiation. Gated for single cells. Representative plot of n = 4 separate cultures. **C**: Quantification of endoderm and hemogenic mesoderm markers on day 3 assessed *via* flow cytometry. n = 4 separate cultures shown as individual data points, error bars min to max. **D**: Assessing effects of including BMP/FGF2 in the FLO-generation protocol on hematopoietic and epithelial marker expression. Representative flow cytometry results on day 18 are shown for n=3 independent experiments. **E**: Spontaneous formation of red blood cell colonies (day 15 top, day 24 middle) and vessel-like structures (day 24 bottom). Representative image of n > 30 separate organoid cultures. Scale bar 100 µm. **F**: 3D-light sheet microscopy of FLO on day 36 after whole-mount staining with fluorescent CD34-antibody, CD31- and ECAD-nanobody. Representative image of n = 4 separate organoid cultures. Scale bar 500 µm. **G**: Surface rendering based on light sheet microscopy image (top) with 3D visualization of CD31+ and CD34+ vessel-like and CD34+ single-cell structures (bottom). Representative image of n = 4 separate organoid cultures. Scale bar 200 µm (top), grid 100 µm (bottom). **H**: RT-qPCR analysis of whole FLO samples for hepatic (*AFP, ALB*) and hematopoietic (*EPO, RUNX1*) markers on different culture days relative to day 6. n=3 separate organoid cultures shown as mean ± SD. FC = Fold Change. **I**: FLO-generation with an ESC(H9C61)-*RUNX1+23-*enhancer*-*DsRed-reporter line demonstrating the temporal relationship of RUNX1-activation to erythroid colony emergence. Representative images shown of n = 3 separate organoid cultures. Scale bar 100 µm. **J**: Multiplexed immunofluorescence (mIF) of FFPE embedded FLOs on day 24 and human fetal liver, 16 weeks post conception (pcw). For FLOs, representative images of n=4 independent organoid cultures are shown. Scale bar 50 µm. **K:** Heat map of RT-qPCR of whole FLOs on day 24, hiPSCs, iPSC-derived 2D hepatocytes (iHeps), primary adult hepatocytes (AH) adult liver sample from explant (liver) for hepatoblast (HB; *FST, APOM, SPINK1, IGF2*), fetal (FH; *AFP, APOA1, GC*) and adult (AH; *ADH1, CYP3A4*) hepatocyte genes according to Wesley et al. (Wesley et al., 2022), normalized *via* dCq to *GAPDH*. Each dataset represents n = 3-5 separate cultures for FLOs and iHeps, n=3 donors of *ex vivo* AHs, and one liver explant.

To assess the developmental stage of the hepatocellular compartment within FLOs, we compared embryonic, fetal, and adult expression modules from genes established in a published Developmental Fetal Liver Atlas (Wesley et al., 2022). We found a fetal-predominant hepatic signature (*AFP, APOA1,* and *GC*) with partial embryonic (*IGF2*) and adult (*CYP3A4, ADH1*) profiles (Fig. 1K, Supplementary Table 2). We further assessed the hepatocyte-specific functionality and examined differences over yolk sac endoderm or published yolk sac organoids (Tamaoki et al., 2023). We conducted cytochrome P450 3A4 (CYP3A4) assay and found 2.84% ± 0.45 (SD) mean activity in whole FLOs relative to primary adult hepatocytes (AH). We contextualized these findings and found that conventional hiPSC-derived fetal-like hepatocyte differentiation protocol (iHeps) in 2D monoculture (Kobayashi et al., 2023) demonstrated 9.53% ± 0.55 (SD) mean activity relative to AHs (Fig. S1E). Further evaluation of the FLO-hepatocellular compartment revealed a gene signature indicative of hepatocytes and not primary or organoid-derived yolk sac endoderm (Goh et al., 2023; Tamaoki et al., 2023), such as liver-specific coagulation factors (*F11, F13b)* and detoxification enzyme *(e.g., ADH1A*; Fig. S1F). Thus, FLOs generated without added hepatocyte-differentiation factors can develop a liver-specific fetal hepatocyte compartment only after receiving initial co-inductive stimuli until day 10.

In human liver development, different levels of hemoglobin subtype representations can help track erythropoiesis to predominantly embryonic yolk sac-like versus embryonic or fetal liver-like (Goh et al., 2023; Peschle et al., 1985). In FLO cultures, we found the predominant presence of a fetal hemoglobin (HbF) signature *(HBG1/2* and *HBA1/2*), albeit a memory of embryonic yolk sac-like zeta (*HBZ*) and epsilon chain (*HBE1*) expression persisted (Fig. S1G, Supplementary Table 2). Therefore, developing an evolving yolk sac-to-liver hematopoietic program within a hepatic niche architecture indicates that FLOs function as an integrated developmental model different from previous models of the yolk sac or the liver alone (Atkins et al., 2022; Hu et al., 2018; Takebe et al., 2013; Tamaoki et al., 2023; Wesley et al., 2022; Zhu et al., 2014).

### FLOs harbor multipotent hematopoietic progenitor cells (HPCs)

We detected CD34+ CD45+ cells in FLOs and fetal liver indicative of HPCs (Fig. 2A). To dissect FLO’s intrinsic from inducible hematopoietic potential, we generated FLOs with exogenously added stimuli. GM-CSF supports macrophages and granulocyte differentiation (Becher et al., 2016). IL-34 is increased during development and may promote tissue-resident macrophage ontogeny (Muñoz-Garcia et al., 2021). UM171 is known to expand HSPCs *in vitro* by restoring histone marks deranged *ex vivo* and promoting myeloid and lymphoid differentiation (Chagraoui et al., 2021; Cohen et al., 2020; Mesquitta et al., 2019). IL-3 is a multilineage colony-stimulating factor (Ackermann et al., 2021; Robin et al., 2006; Umemura et al., 1989). We included this small molecule-cytokine cocktail (“GM/34”) as an additional experimental arm followed by CD34+ isolation for immunophenotypical and functional hematopoietic characterization (Fig. 2B).

**Figure 2.**
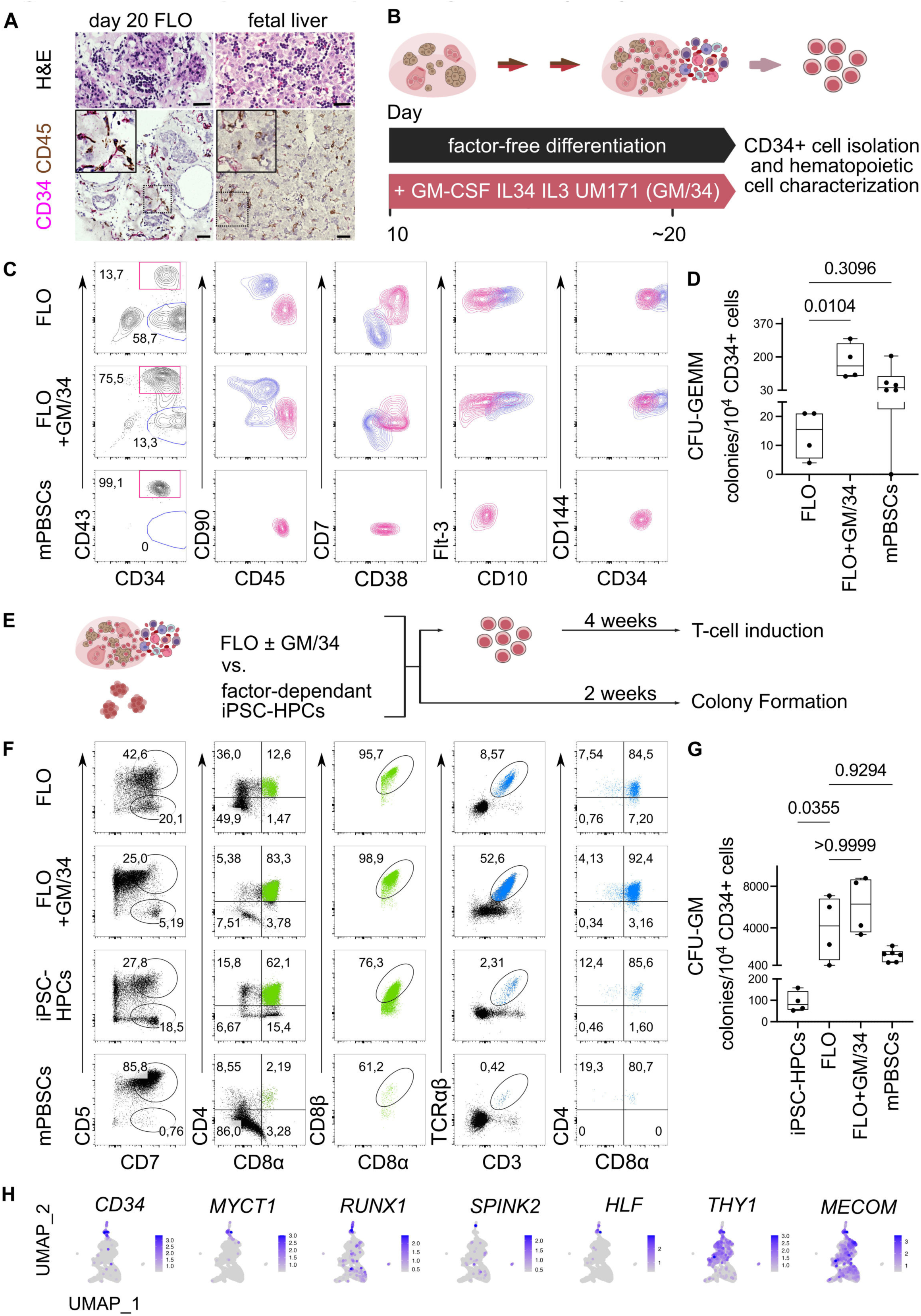
FLOs harbor multipotent Hematopoietic Progenitor Cells (HPCs) **A**: H&E and IHC for indicated markers on day 20-FLOs and fetal liver 18 pcw. Scale bar 50 µm. **B**: Schematic of CD34+ HPC isolation for hematopoietic characterization from factor-free differentiated FLOs and FLOs with factors GM-CSF, IL34, IL3, and UM171 or UM729 (“FLO + GM/34”) added from day 10. **C**: Flow cytometry analysis of MACS-CD34 enriched FLO, FLO + GM/34 on days 18-22, and mobilized peripheral blood stem cells (mPBSCs). Gated for viable single cells. The blue population is gated for CD34+ CD43-. The pink population is gated for CD34+ CD43+. Pooled experiment of 40 separate organoid cultures. **D**: Granulocyte, erythroid, macrophage, and megakaryocyte colony forming units (CFU-GEMM) after two weeks culture in MethoCult SF H4436 (STEMCELL Technologies). FLOs and FLOs + GM/34 on days 22-27 were CD34+ enriched *via* MACS before plating. Primary CD34+ mPBSCs (2 healthy donors) are shown as controls. The colony numbers were normalized to 10000 plated CD34+ cells assessed *via* flow cytometry. n = 4 (for FLO, FLO + GM/34) and n = 6 (for mPBSCs); separate cultures are shown as individual data points min to max. Kruskal-Wallis with Dunńs multiple comparison correction was used to assess significance. Corrected p-values shown. **E**: Schematic of functional hematopoietic differentiation assays of FLOs with or without GM/34 compared to adapted factor-dependent iPSC differentiation protocol of HPCs (iPCS-HPCs). **F**: Flow cytometry analysis for T-cell markers after four weeks of culture using StemDiff T Cell Kit (STEMCELL Technologies). CD34+ cells were enriched *via* MACS from FLOs, FLOs + GM/34 on days 18-22 and iPSC-HPCs before plating. Primary human mPBSCs (healthy donor) were used as a control. Gated for viable single cells. Pooled data of n = 40 separate organoid cultures. **G**: Granulocyte-macrophage colony forming units (CFU-GM) after 2 weeks culture in MethoCult SF H4436 of FLOs and FLOs + GM/34 on days 22-27. Primary CD34+ mPBSCs (2 healthy donors) and iPSC-HPCs are shown as controls. The colony numbers were normalized 10000 plated CD34+ cells assessed *via* flow cytometry. n = 4 (for FLO, FLO + GM/34 and iPSC-HPCs) and n = 6 (for mPBSCs); separate cultures are shown as individual data points min to max. Kruskal-Wallis with Dunńs multiple comparison correction was used to assess significance. Corrected p-values are shown. **H:** UMAP feature plots of FLOs + GM/34 scRNA-seq datasets on day 13 displaying expression levels of hematopoietic stem and progenitor (HSPC) signature genes.

We performed spectral flow cytometry and compared the results with primary human CD34+ mobilized peripheral blood stem cells (mPBSCs). Without added hematopoietic cytokines, CD34+ enriched FLOs contained 13.7% CD34+ CD43+ cells. The addition of GM/34 increased the frequency by almost 6-fold to 75.5% (Fig. 2C). The immunophenotype within the CD34+CD43+ subset was predominantly negative for CD90 and CD10, partially negative for CD38 and CD7, and positive for CD45 and FLT3 consistent with reports of multipotent HPCs (Doulatov et al., 2010). This immunophenotype resembled mPBSC, albeit the primary control cells displayed less cellular heterogeneity overall. CD144 has been reported to be expressed in murine fetal hematopoiesis but not the adult counterparts (Kim et al., 2005). FLOs-HPCs retained the endothelial marker CD144, unlike the adult HSC control (Fig. 2C). To validate multipotency with and without added cytokines, CD34+ enriched cells of each arm were subjected to colony-forming units (CFU)-assays. Within two weeks, granulocyte, erythroid, macrophage and megakaryocyte colony-forming units (CFU-GEMM) emerged from FLO cultures. Adding GM/34 significantly increased the colony-forming efficiency of FLOs by about 10-fold to levels observed with primary HSCs (Fig. 2D, S2A). Thus, FLOs at baseline can self-develop, mature, and harbor functional erythromyeloid progenitors, whose CFU efficiency can be augmented by adding GM/34.

To dissect the inherent lymphoid potential, we exposed FLO-derived HPCs to a T-cell differentiation assay and compared the results with those obtained from state-of-the-art iPSC-derived HPCs (iPSC-HPCs) from the same iPSC-line (Fig 2E). After four weeks of T-cell induction, we observed a CD5+ CD7+ population highly abundant in mPBSCs (85.8%), which emerges first in T-cell development (Seet et al., 2017) (Fig. 2F). CD4+ CD8α+ double positive (DP) cells, indicative of T precursor cells, were detected in FLOs (12.6%) and drastically increased in FLO + GM/34 (83.3%), with over 95% of the DP population co-expressing CD8β. Remarkably, 8.6% and 53.6% of the cells expressed CD3+ TCRαβ+ in FLOs and FLOs + GM/34, respectively, exceeding the levels observed in iPSC-HPCs (2.31%) and mPBSCs (0.42%). Interestingly, a small subset of CD3+ TCRαβ+ CD4+ CD8α-so-called single-positive cells (SP) indicated some degree of maturation (Fig. 2F). These findings indicate that FLOs harbor lymphoid potential. Further, this lymphoid potential appears more mature at baseline than those differentiated from iPSC-HPCs or mPBSCs and is tractable by adding cytokines. We further confirmed multipotency by exposing FLOs to MS5 cell co-culture to assess B-cell potential. We exhibited a CD45+ CD20+ CD43+ CD27+ B cell immunophenotype, similar to human B1 cells in the human fetal tissues and cord blood (Griffin et al., 2011; Suo et al., 2022) (Fig. S2B). To support the hypothesis that the FLO microenvironment is favorable for multi-lineage hematopoiesis, we assessed the myeloid potential of whole cultures while preserving the non-hematopoietic cells. FLOs generated significantly more granulocyte, macrophage CFUs (CFU-GMs) than iPSC-HPCs lacking the FLO microenvironment (Fig. 2G). FLOs colony output was unchanged from GM/34 or mPBSCs. Thus, FLOs created a self-sufficient hepatic microenvironment supportive of multipotent HPCs, yielding at baseline higher levels of myeloid output relative to iPSC-derived HPCs lacking a hepatic context.

Next, we performed scRNA-Seq on FLO + GM/34 and benchmarked integrated data sets from day 10, 13, 16 and 20 to the published nascent HSC signature scorecard (Calvanese et al., 2022). In our integrated data, we identified, through more resolved clustering, a small hematopoietic subset that expressed most of the “HSC-enriched” gene module but not the “HSC” module (Fig. S2C, D). Analysis of CD34+ cells from FLOs versus human fetal liver (FL) and yolk sac (YS) reveal a yolk sac-like activation of hematopoietic transcription factors and hemogenic endothelial gene programs on days 10 and 13. From day 16, the hematopoietic signature in FLOs increasingly resembled the native fetal liver (Wang et al., 2021). This switch is exemplified by the surge of *BCL11A, PRSS57, SPINK2, HOPX* expression from day 16 (Fig. S2E). HSC-signature genes *CD34, MYCT1, RUNX1, SPINK2, HLF, THY1* and *MECOM* were co-expressed on day 13 (Fig. 2H). Further, marker genes of endothelial–to–hematopoietic transition (EHT) were co-expressed on day 13, which diverged into distinct areas on day 16 (Fig. S2F). This observation supports the concept that FLO development consists of two stages: an early establishment phase between days 10 and 13, coinciding with EHT and FLO assembly, and a second fetal liver-like hematopoietic stage from day 16.

### FLOs establish a functional hematopoietic niche, including hepatic-to-hematopoietic crosstalk

We characterized and validated the intrinsic FLO niche signaling across cell compartments. We observed co-localization of HepPar1+ hepatocyte-like cells with CXCL12, a critical chemoattractant regulating mobilization and homing of HSPCs in the fetal liver and bone marrow (Lapidot et al., 2005; Soares-da-Silva et al., 2020). Elongated and round CD34+ cells co-expressed its receptor CXCR4 with a partial phosphorylation pattern, indicating variable CXCL12/CXCR4 axis activation (Fig. 3A). CD34+ CXCR4+ cells spatially clustered significantly in closer proximity to HepPar1+ when co-expressing CXCL12 (Fig. S3A, B). In FLOs, the secreted CXCL12 concentration increased over time (Fig. 3B, S3C). A migration assay, with and without receptor inhibition (AMD3100, AMD), demonstrated that primary CD34+ cells migrated towards a FLO-derived CXCL12 gradient using a transwell culture per CXCR4-specific chemoattraction (Fig. 3C, D). Further, when adding the same CXCR4 inhibitor AMD3100 to FLO cultures from day 26 for two weeks, we observed depletion of CD45+ leukocytes in FLOs (Fig. S3D). These indicate that the CXCL12/CXCR4 axis is an intercellular signaling pathway governing migration and leukopoiesis in FLOs.

**Figure 3.**
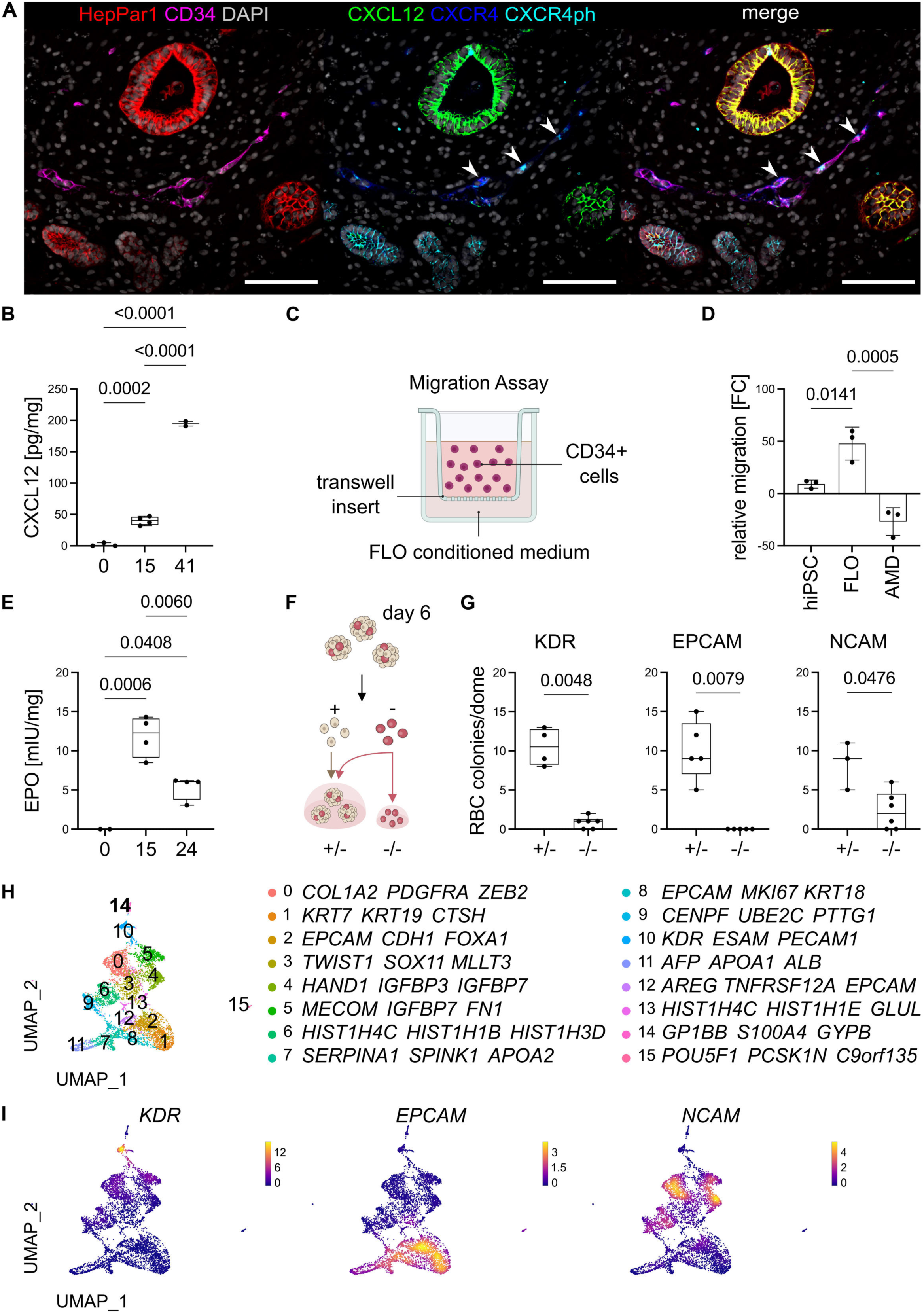
FLOs establish a Hematopoietic Niche with Functional Hepatocyte-to-Hematopoietic Cell Crosstalk. **A**: mIF image of FLOs on day 34 demonstrating co-localization of HepPar1 with CXCL12, CD34 with CXCR4, and CXCR4 phosphorylation (CXCR4ph) of CD34+ cells (indicated by arrows). Representative image of n = 8 separate organoid cultures. Scale bar 100 µm. **B**: Protein concentration of CXCL12 on days 0, 15, and 42 in FLO-conditioned medium assessed *via* ELISA. Results are normalized to 1 mg total protein per 48 hours. n = 2-4 separate organoid cultures are shown as individual data points min to max. One-way ANOVA with Tukey multiple comparison test was used to test for significance. Corrected p-values are shown. **C**: Schematic of migration assay using primary human CD34+ cells. Created with BioRender.com. **D**: CXCL12-dependent chemoattraction shown by migration of primary human CD34+ cells from mPBSCs in conditioned medium from hiPSCs, FLOs, and FLOs with CXCR4 inhibition using AMD3100 (AMD). n = 3 separate cultures shown as individual data points and mean ± SD. One-way ANOVA with Dunnett multiple comparison test to assess significance. Corrected p-values are shown. **E**: Protein concentration of EPO on days 0, 15, and 24 in FLO-conditioned medium assessed *via* ELISA. Results are normalized to 1 mg total protein per 48 hours. n = 2-4 separate organoid cultures are shown as individual data points min to max. One-way ANOVA with Tukey multiple comparison test was used to test for significance. Corrected p-values are shown. **F**: Schematic of the lineage depletion approach using FACS-sorting on day 6 for respective markers followed by embedding in Matrigel domes as isolated negative populations (−/−) or reconstituted with respective positive populations (+/−) according to the initial cell ratio. **G**: Quantification of red blood cell (RBC) colonies per dome on day 15 organoids reconstituted (+/−) or depleted (−/−) for KDR (left), EPCAM (center), and NCAM (right). n = 3-6 separate organoid cultures are shown as individual data points min to max. Two-way ANOVA with Sidak’s multiple comparison test was used to assess significance. Corrected p-values are shown. **H**: UMAP of Seurat-guided clustering of FLO + GM/34 scRNA-seq dataset on day 13. Representative genes of the top 20 expressed genes relative to all clusters are shown for each cluster (Supplementary Table 3). **I**: UMAP feature plots of FLOs + GM/34 scRNA-seq datasets on day 13 for *KDR*, *EPCAM,* and *NCAM*.

We detected secreted EPO in the FLO microenvironment with a peak on day 15 (Fig. 3E). To test how lineage crosstalk promotes erythropoiesis, we fractionated FLO cultures on day 6 for EPCAM (marking fetal hepatocytes and epithelium), KDR (marking hemogenic endothelial cells and mesenchyme) and NCAM (marking mesenchyme) followed by seeding in Matrigel domes as isolated negative populations or reconstituted with respective positive populations. The loss of epithelium completely depleted erythropoiesis. Interestingly, also the loss of KDR+ and NCAM+ compartments reached statistical significance in compromising red blood cell output (Fig. 1F, G). scRNA-Seq data sets of day 13 FLOs identified emerging clusters representing progeny from endothelial (cluster 10), epithelial (clusters 1, 2, 7, 8, 11, 12) and mesenchymal (clusters 0, 3, 4, 5) precursors co-expressing *KDR, EPCAM* and *NCAM,* respectively (Fig. 3H, I, Supplementary Table 3). Thus, FLOs induce erythropoiesis through EPO secretion which emerges *via* endothelial, epithelial, and mesenchymal lineage crosstalk.

To further validate the impact of FLO-niche cells, we performed a CFU assay comparing FLO stromal-reduced cultures (CD34 enriched) to whole FLO cultures, where the FLO stroma was preserved. We observed more CFU-GM colonies when preserving the FLO stroma (Fig. S3E). Macrophage colony-stimulating factor (M-CSF) was detected in FLO cultures (Fig. S3F). Blocking the CSF-1 receptor with BLZ945 (BLZ) blocked most immune cell differentiation, indicating that the FLO microenvironment supports myelopoiesis (Fig. S3G).

scRNA-Seq data sets of day 20 FLOs mapped a set of gene ontology biological process (GOBP) genes for positive regulation of hematopoietic progenitor cell differentiation–that includes *KITLG*–to mostly mesenchyme-, but also to epithelial-associated clusters, especially the cholangiocyte gene-expressing cluster 13 (Fig. S4A, B, Supplementary Table 4). We tracked the expression of *EPO* specifically to cluster 12 – predominantly expressing hepatocyte genes. We found the same expression pattern for the hematopoietic niche factors *IGF2, ANGPTL2,* and *ANGPTL3*. *CSF-1* and pericyte/mesenchyme marker *NES* are co-expressed (Fig. S4C). On the protein level, we confirmed the presence of IGF2+ epithelium and Nestin+, which are known components of the fetal liver microenvironment (Fig S4D, E) (Soares-da-Silva et al., 2020). These findings show that FLOs develop a complex hematopoietic niche tissue with mesenchymal and hepatobiliary contribution that resembles aspects of hepatic ontogeny.

### Longitudinal analysis of multi-lineage trajectories and myeloid diversity

After integrating our scRNA-seq data set from FLO + GM/34 on days 10, 13, 16, and 20, we identified a distinct hematopoietic population with multiple trajectories (expressing *CD34, RUNX1, HLF, SPINK2*), that was inferred to be separate from the endoderm-to-hepatocyte-lineage trajectory (*PROX1, FOXA2, AFP, ALB, HNF4A, CYP3A4*; Fig. 4A). Inferred progenitor activities *via* CytoTrace (Gulati et al., 2020) suggested progenitor activity in a subset of hematopoietic cells (Fig. S5A). Annotation to Human Fetal Liver reference atlas established similarities to most hematopoietic and immune cell states (21 of 24, including multiple myeloid progenitors, e.g. neutrophil-myeloid progenitors and a smaller subset of lymphoid precursors), and all non-immune cell states (3 of 3) (Fig. S5B) (Popescu et al., 2019). We also integrated FLOs into an independent Fetal Liver (FL) and Yolk Sac (YS) dataset and compared both types of annotations. FLOs recapitulated most cell states, albeit megakaryocytes, erythroid, and mast cells were underrepresented, while myeloid progenitors and progeny were highly represented (Fig. S5C-E) (Wang et al., 2021). In our comparison of FLOs with human postnatal liver using mIF, we verified the presence of HepPar1+ or CK19+ hepatobiliary, PDGFRA+ mesenchymal, and IBA1+ immune cells. Of note, the presence of MPO+ cells in FLOs required the addition of GM/34 (Fig. 4B). This finding is consistent with the notion that neutrophil-myeloid progenitors exist in the human fetal liver (Popescu et al., 2019), but may need extrahepatic granulocytic signals for differentiation. IHC confirmed additional granulocyte states in addition to myeloid or macrophage states as evidenced by stainings for neutrophil elastase (NE) in addition to CD68 and general myeloid cells (CD15) (Fig. S5F). To understand the myeloid progenitor landscape in FLOs, we looked for hotspots within the hematopoietic scRNASeq population expressing published gene sets suggestive of specific progenitor states. We found distinct signatures indicating HSPC-like (*RUNX1, HOXA9, MECOM, HLF, MLLT3, SPINK2*) (Calvanese et al., 2022), yolk sac-derived myeloid-biased progenitor-like (YSMP; *CD34, KIT, MYB, AZU1, MPO, MS4A3, LYZ*), YSMP-derived embryonic monocyte-like (YSMP Mono; *CCR2, HLA-DR, CSF1R, MEF2C, CD14*) and YSMP-derived embryonic granulocyte-like (YSMP Granulo; *S100P, S100A8, S100A9, S100A12, CEACAM8/*CD66b) hotspots (Fig. 4C)(Bian et al., 2020). While expression patterns in clusters 6 and 7 resemble YSMPs, FLOs further demonstrated gene expression profiles of neutrophilic (cluster 1) and monocytic (clusters 5 and 8) differentiation trajectories in the embryonic liver (Fig. 4D, E, S5G). Next, we assessed whether FLOs can differentiate further along transcriptionally inferred trajectories. We added G-CSF from day 20 to our FLO + GM/34 cultures to further mature granulocyte differentiation and performed FACS on day 30 for immunophenotype characterization. Indeed, we identified distinct CD45+ CD14-CD16+ CD66b+ neutrophil granulocyte and CD45+ CD14+ CD163+ macrophage populations (Fig. 4F, G). Giemsa staining of respective isolated populations confirmed neutrophil or histocyte macrophage phenotypes (Fig. 4H). Thus, FLOs harbor a heterogeneous progenitor population, which gives rise to multiple myeloid lineages.

**Figure 4.**
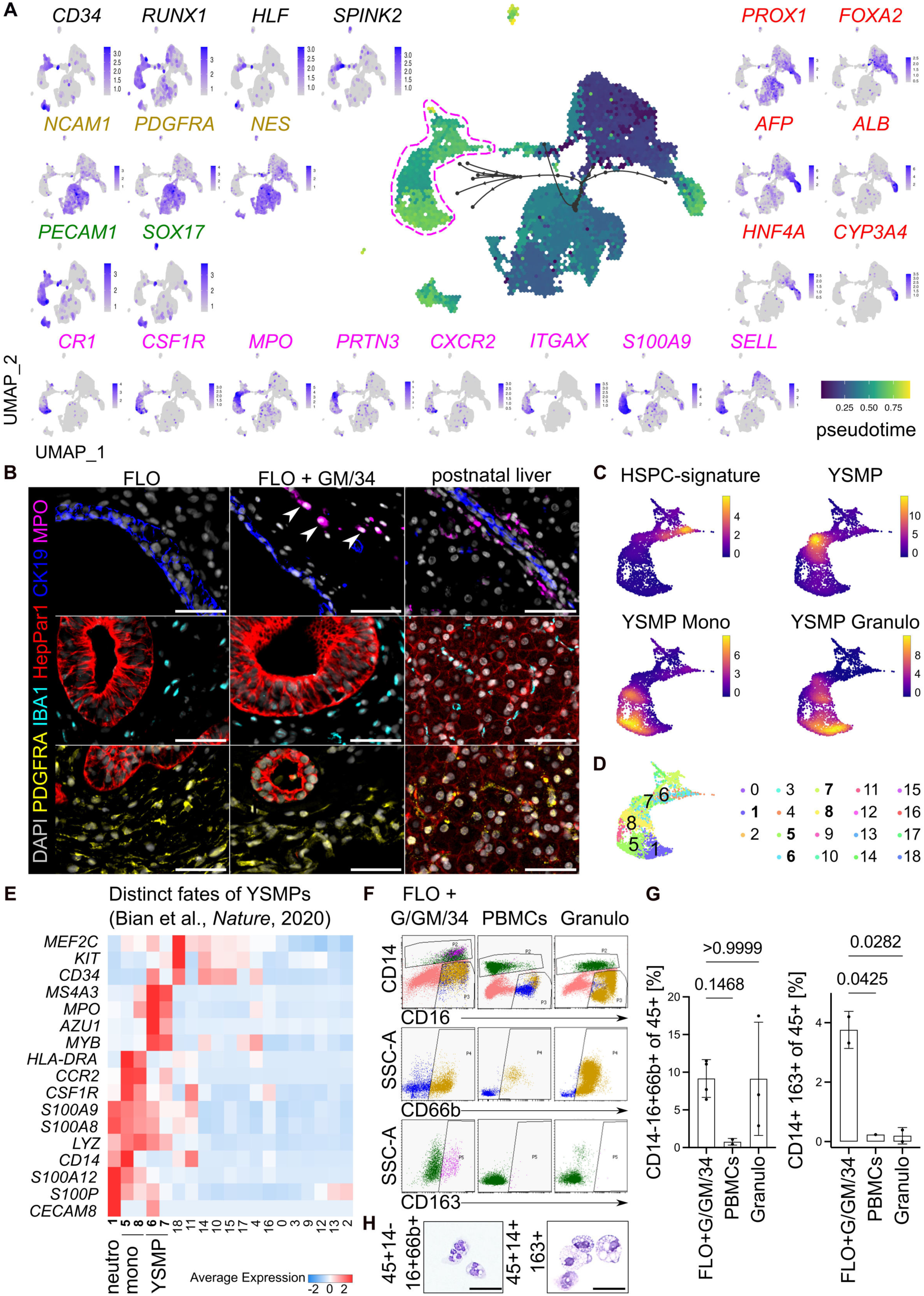
Longitudinal Analysis of FLO-Development and Myeloid Diversity. **A:** UMAP of integrated scRNA-seq datasets from FLOs +GM/34 on days 10, 13, 16, and 20 with pseudo-time (color-coded) and trajectory analysis using PAGA-Tree (indicated in black arrows, hematopoietic cluster highlighted in magenta dotted lines). Feature gene expression plots for maker genes of HPCs (black), stromal mesenchyme (yellow), endothelium (green), myeloid lineages (magenta), endoderm and hepatoblasts/fetal hepatocytes (red). **B:** mIF image of day 34 FLO and FLO + GM/34 compared to postnatal liver specimen (24 months) to assess cellular heterogeneity and the emergence of putative neutrophils (MPO). Representative images of n = 4 separate organoid cultures. Scale bar 50 µm. **C**: UMAP of integrated FLO + GM/34 scRNA-seq datasets (days 10, 13, 16, 20) with a close-up of the hematopoietic cluster. Featuring the gene set expression of HSPC-signature: *RUNX1, HOXA9, MECOM, HLF, MLLT3, SPINK2* from Calvanese et al., *Nature*, 2020(Calvanese et al., 2022); yolk sac-derived myeloid-biased progenitors (YSMPs): *CD34, KIT, MYB, AZU1, MPO, MS4A3, LYZ*; YSMP-derived embryonic monocyte lineage (YSMP Mono): *CCR2, HLA-DR, CSF1R, MEF2C, CD14;* and YSMP-derived embryonic granulocyte lineage (YSMP Granulo): *S100P, S100A8, S100A9, S100A12, CEACAM8/*CD66b from Bian et al. (Bian et al., 2020). Illustrated with https://github.com/ZornLab/Mona. **D**: UMAP demonstrating cell clusters (RNA_snn_res.0.8) of integrated FLO + GM/34 scRNA-seq datasets (days 10, 13, 16, 20) with close-up on the hematopoietic cluster. **E**: Cluster gene expression from integrated FLO + GM/34 dataset (days 10, 13, 16, 20) of markers for YSMP, neutrophilic (neutro) and monocytic (mono) fates per Bian et al. (Bian et al., 2020). **F**: FACS gating strategy to characterize neutrophil granulocytes (CD45+ CD14-CD16+ CD66b+, yellow) and macrophages (CD45+ CD14+ CD163+, magenta) of day 30 FLOs + G/GM/34 (added G-CSF from day 20 to 30 to FLO + GM/34 protocol) compared to fresh adult PBMCs and PBMC-derived granulocytes (Granulo, healthy donor). Gated for viable single cells and CD45+. **G**: Quantification of CD45+ CD14-CD16+ CD66b+ neutrophil granulocytes (left) and CD45+ CD14+ CD163+ macrophages (right) in FLO + G/GM/34 compared to PBMCs and granulocytes (Granulo). n = 2-3 separate organoid cultures shown as individual data points and mean ± SD. One-way ANOVA with Dunnett multiple comparison test to assess significance. Corrected p-values are shown. **H:** Representative images of Giemsa-staining after FACS-based isolation for CD45+ CD14-CD16+ CD66b+ neutrophil granulocytes (left) and CD45+ CD14+ CD163+ macrophages (right). Scale bar 50 µm.

### In-depth characterization and functional assessment of macrophages and granulocytes

Yolk sac-derived macrophages can follow a monocyte or a pre-macrophage trajectory, albeit liver macrophages are considered to derive from embryonic monocytes relying on known myeloid transcription factors (TFs) such as *IRF8, CEBPA,* and *SPI1 (Goh et al., 2023)*. After confirming the expression of this TF-combination in our scRNA-Seq dataset (data not shown), we fractionated FLO + GM/34 for CD14+ CD16+ immune cells and performed bulk RNA sequencing. These CD14+CD16+ cells did not resemble adult classical, nonclassical, or intermediate monocytes but anti-inflammatory-polarized macrophage controls. Similar hierarchical clustering patterns were obtained regardless of filtering for human monocyte gene panels, CD163-subset-isolation, or lipotoxic injury (Fig. 5A, S6A). scRNA-seq data sets from day 20 FLO + GM/34 revealed a subset of cells with a Kupffer-like signature, namely *VSIG4, MARCO, ID1, CD14, CD163, CD68, FCG3A* (Fig. 5B). Flow cytometry analysis of FLOs on day 30 revealed a predominant CD45+ CD64low population, indicative of an anti-inflammatory macrophage phenotype, but lacking CD206 expression. The addition of GM/34 during FLO differentiation further enriched this CD45+ CD64low population and induced CD206 expression (Fig. 5C). Thus, we observed in FLOs the developmental handover from embryonic monocyte to hepatic macrophage trajectory and that GM/34 induced in-organoid macrophage maturation.

**Figure 5.**
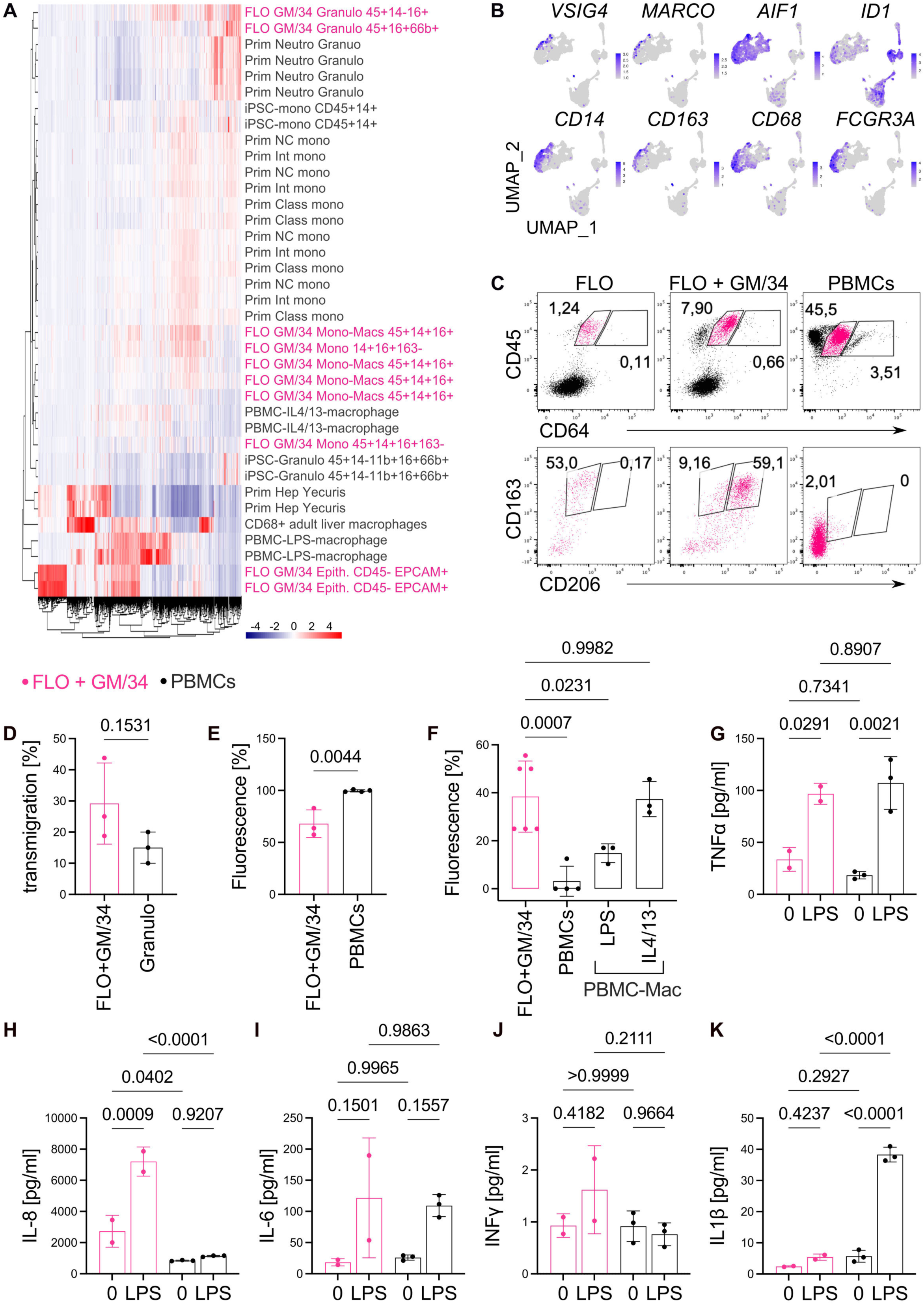
Functional Myeloid Characterization and Macrophage Polarization. **A:** Gene expression of bulk RNA-seq data with unbiased hierarchical clustering analysis of primary cells, FLO + GM/34-derived myeloid subsets, and other iPSC-derived immune cells. FLO + GM/34 derivatives are highlighted in magenta. The primary immune cells include LPS and IL4/IL13-polarized PBMC-derived macrophages, neutrophil granulocytes (Prim Neutro Granulo), monocytes (Class=classical, Int=intermediate, and NC=nonclassical subsets, monocyte data derived from Monaco et al(Monaco et al., 2019). and integrated into our dataset), and CD68+ adult liver macrophages. iPSC-derived neutrophil granulocytes and -monocyte differentiated as per Trump et al. (Trump et al., 2019)(iPSC-Neutro Granulo and -Mono, respectively), and primary hepatocytes (Prim Hep Yecuris) were included. Each sample represents a separate culture. **B**: UMAP feature plots of FLOs + GM/34 scRNA-seq dataset on day 20 displaying expression levels of Kupffer cell signature genes. **C**: Anti-inflammatory polarization of macrophages in FLOs + GM/34 shown by CD45+ CD64low CD162+ CD206+ compared to FLOs without GM/34 and primary PBMCs (healthy donor) assessed *via* flow cytometry. Gated for viable single cells, and for CD45+ CD64low (magenta). Representative plots of n = 2 separate cultures. **D**: Granulocyte functional assay showing percentage of granulocytes (seeded in a transwell) that migrated towards an fMLP-gradient of FLO + GM/34-derived granulocytes (CD45+ CD16+ CD66b+ FACS sorted) and primary human granulocytes (Granulo). n = 3 separate organoid cultures (for FLO + GM/34) and n = 3 technical replicates (for Granulo) shown as individual data points and mean ± SD. An unpaired two-tailed t-test was used to assess significance. p-values are shown. **E**: DCFH-DiOxyQ probe-based fluorescent detection of hydrogen peroxide (H_2_O_2_), peroxyl radical (ROO·), nitric oxide (NO), and peroxynitrite anion (ONOO-) of FLO + GM/34-derived granulocytes (CD45+ CD16+ CD66b+ FACS sorted) and PBMCs. n = 3 separate organoid cultures (for FLO + GM/34) and n = 4 technical replicates (for PBMCs, single donor) shown as individual data points and mean ± SD. An unpaired two-tailed t-test was used to assess significance. p-values are shown. **F**: Phagocytosis-activity of freshly thawed PBMCs, PBMC-derived macrophages (LPS or IL4/IL13-polarized), and FLO + GM/34-derived monocyte-macrophages (CD45+ CD14+ CD16+ FACS sorted) after exposure to pHrodo™ Deep Red E. coli BioParticles™. n = 6 separate organoid cultures shown as individual data points and mean ± SD. Ordinary one-way ANOVA with Dunnett multiple comparisons test was used to test for significance. Corrected p-values are shown. **G-K**: Multiplex cytokine detection assay for FACS-sorted CD14+CD16+ cells (10000 cells per sample) from PBMCs (black) and FLOs + GM/34 (magenta). Cells were either untreated (0) or LPS treated (LPS). n = 2 separate organoid cultures (for FLO + GM/34) and n = 3 technical replicates (for PBMCs) shown as individual data points and mean ± SD. Two-way ANOVA with Tukey’s multiple comparisons test was used to test significance. Corrected p-values are shown for 0 vs. LPS of respective samples and PBMCs vs. FLOs of respective conditions.

To evaluate the functionality of FLO + GM/34-derived granulocytes and macrophages, we conducted FACS sorting of respective CD45+ CD16+ CD66b+ versus CD45+ CD14+ CD16+ populations followed by assays testing for granulocyte migration, reactive oxidative species formation, phagocytosis, and cytokine secretion upon lipotoxic exposure. Sorted CD45+ CD16+ CD66b+ granulocytes migrated towards a chemokine gradient similar to adult granulocytes, though a reduced level of reactive oxidative species formation (H_2_O_2_, ROO·, NO, ONOO-) was observed (Fig. 5D, E). Sorted CD45+14+16+ FLO-macrophages exhibited high phagocytic activity to bacterial particles, comparable to adult PBMC-monocyte-derived IL4/IL13-polarized primary macrophages (Fig. 5F). Moreover, FLO + GM/34 macrophages displayed a cytokine secretion profile resembling postnatal PBMC-derived controls and demonstrating elevated sensitivity for IL-8 upon endotoxin stimulation (Fig. 5G-K). These findings suggest that FLO + GM/34-derived macrophages are immune-reactive and respond to bacterial stimuli.

We transplanted whole FLOs generated with *RUNX1* +23 enhancer dsRed enhancer reporter ESC line into the kidney capsule (KC) of NSG mice. After two weeks, the entire FLO microenvironment engrafted, as evidenced by the presence of human (NM95+) AFP+, CD34+, CK19+, IBA1+ and CD31+ cells (Fig. S6B-D). We investigated hematopoietic activity *in vivo* by mpIF and detected human CD34+ hCD45+ cells, and *RUNX1* enhancer reporter signal using RFP-antibody against dsRed (Fig. S6E). Flow cytometry of transplanted whole kidneys detected 0.2% viable mCD45-hCD45+ hCD34+ cells after two weeks (Fig. S6F). After six weeks, we observed NM95+ CD34+ CD31+ capillary endothelial-like structures within the transplanted area (Fig. S6H). The migration of FLO-derived myeloid cells into other tissues was examined using flow cytometry. After six weeks, 1.50% of mCD45+ hCD45+ CD16/32+ cells were detected in the mouse liver but not in other organs (Fig. S6I and data not shown). Taken together, FLOs establish an ectopic hematopoietic tissue *in vivo*, and hepatic ontogeny of FLO-myeloid cells may result in liver-biased migration reminiscent of hepatic Kupffer cell engraftment.

## DISCUSSION

Current models of the human fetal liver do not recapitulate the hematopoietic complexity found in human ontogeny (Goh et al., 2023; Popescu et al., 2019; Vanuytsel et al., 2022; Wang et al., 2021; Wesley et al., 2022). The inability to mechanistically interrogate the multicellular fetal liver niche limits understanding human development and translational studies of infant leukemias, immune deficiencies, and inflammatory hepatopathies.

To investigate human fetal liver tissue, we developed FLOs as a miniature model of the tissue. These FLOs are designed to mimic the environment of the fetal liver and include a yolk sac-like hematopoietic system. Specifically, hematopoietic benchmarks in FLOs resemble a yolk sac-derived wave of lineage-biased HPCs with erythromyeloid and retained lymphoid potential (Canu & Ruhrberg, 2021; Goh et al., 2023).

Interestingly, the *HBG1*-predominant globin signature confirmed a fetal liver-like program, albeit the embryonic hemoglobin memory suggests progenitor programs in developing FLOs *per se* may be heterogeneous and recapitulate different waves. Because direct evidence has not been established yet that human liver erythropoiesis is mostly sourced from AGM-derived HSPCs rather than YS-derived progenitors, the observation in FLOs supports the hypothesis of YS-derived fetal erythropoiesis.

Although recent progress has been evident in the field of modeling different hematopoietic tissues *in vitro* using hPSCs, e.g., yolk sac hematopoiesis (Atkins et al., 2022), bone marrow-like organoids (Frenz-Wiessner et al., 2024) and AGM-like HSCs that engraft in immune-deficient mice (Ng et al., 2024). FLOs are the first to integrate liver-specific ontogeny into hematopoietic systems, including fetal hepatocytes and mesenchymal cells. We demonstrate that initial germ layer co-induction and signaling until day 10 is sufficient for the hepatocellular compartment to develop. We speculate that niche signaling, e.g., Oncostatin signaling, expressed in FLOs, is harnessed as niche signaling for self-sufficient hepatocyte differentiation, albeit future studies would require specific inhibition studies.

Using ten pluripotent stem cell lines, we established FLOs that resemble the cellular composition of the fetal liver, containing various blood cell types, liver cells, and supporting cells. Our analysis revealed that most native cell types were accurately reproduced in the FLOs. However, we did observe an overrepresentation of myeloid and mesenchymal cells and a lower representation of lymphoid and erythroid cells. Yet, it is important to consider that the methods used to isolate the cells and potential single-cell-genomic artifacts may have impacted these observations.

FLOs recapitulate liver embryonic monocytes previously described by Goh et al. (Goh et al., 2023), which helps to further differentiate FLO hematopoiesis from the early yolk sac primitive hematopoiesis. Assessing the functional maturation of immune cell types in multilineage organoids in detail is challenging. Here, we extensively examined through various methods, including single-cell and bulk RNA sequencing followed by functional testing of multiple myeloid lineages after in-organoid differentiation. We have identified the establishment of embryonic-monocyte-derived functional macrophages with immature polarization, which could be completed with the GM/34 cocktail with an emerging Kupffer-like subset signature.

Our findings demonstrate that FLOs serve as a novel model system facilitating the functional analysis of complex hepato-hematopoietic crosstalk and embryonic-to-early fetal hematopoiesis. We have characterized the heterogeneity of the HPC and progeny population within FLOs, revealing multipotency biased towards myeloid lineage commitment while retaining fetal B and T cell potential. Further studies are required to elucidate the function of *in vivo* engraftment. Preliminary data further suggest that FLO-derived cells exhibit preferential liver engraftment. This raises hope that our approach will be an effective liver-targeted and personalized cell therapy, e.g., to replace hyperinflammatory bone-marrow-derived liver macrophages. Finally, FLOs provide a platform to understand liver organogenesis and warrant further investigation in developmental pathogenesis.

## LIMITATIONS OF THE STUDY

The low representation of fetal hepatocytes in our transcriptional map matches results from human fetal liver maps (Popescu et al., 2019). Still, it may also be related to technical difficulties in single-cell isolation and genomics of hepatocytes. The lymphoid-biased cell states, however, were underrepresented relative to primary liver datasets. This aligns with the well-known challenges of generating lymphocytes *in vitro*. Despite a myelopoietic growth advantage, we could identify multiple lymphopoietic progenitors, which we validated per T and B-cell induction assays. Finally, our experiments were not designed to assess definitive fetal hematopoiesis and focused on determining capacity and requirements for HSC-independent hematopoiesis like early fetal–but not primitive– HSPCs from the Yolk Sac populating the liver. Despite the mentioned limitations, the lymphoid potential in FLOs underscores hematopoietic multipotency. It opens opportunities for future studies involving the co-differentiation of lymphocyte subsets in a hepatobiliary context.

---

**Supplementary Figure S1**

**A**: Zinc finger nucleases-mediated integration of the RUNX1 murine +23 donor dsRed reporter cassette *via* homologous recombination in H9 embryonic stem cells. **B**: Giemsa stain of various identified nucleated erythroid progenitors after CD235a+ sorting on day 18. Scale bar 25 µm. **C**: Percentages of cell compartments in FLOs over time assessed *via* mIF images. Number of positive cells normalized to DAPI. n = 2-4 separate cultures shown as individual data points min to max. **D**: IHC for indicated markers on day 20-FLOs and fetal liver on 39 pcw for AFP, and 18 pcw for ALB. Scale bar 50 µm. **E**: CYP3A4 live cell function assay on primary adult hepatocytes (AH, 24h after thawing onto collagen-coated tissue plates) and day 30 FLOs (left) and conventional PSC-derived hepatocyte differentiation protocol (Kajiwara et al., 2012) (iHeps, right). Results normalized to viable cell number and relative to AH (100%). n=2-4 separate cultures shown as individual data points min to max. An unpaired two-tailed t-test was used to assess significance. p-values are shown. **F:** Heat map of relative expression of integrated FLO + GM/34 scRNA-seq datasets (days 10, 13, 16, 20) featuring genes that are generic for yolk sac endoderm and hepatocyte or specific to hepatocytes (*ADH1, ADH6, CYP3A7, GC*, *F9, F11, F13b, CPB2*) per Goh et al., (Goh et al., 2023). Cell types were predicted using Random Forrest analysis using data from Popescu et al. (Popescu et al., 2019). Hep = hepatocytes, lympho/T = lymphoid/ T lymphocyte, pre = precursor, Ery = erythroid, Ery Island Mac = erythroblastic island macrophage, pro B = pro B cell, mono = monocyte, Neu-my pro = neutrophil-myeloid progenitor, Endo = endothelial cell, Fib = fibroblast, MEMP = megakaryocyte-erythroid-mast cell progenitor, MK = megakaryocyte. Illustrated with https://github.com/ZornLab/Mona. **G**: Heat map of RT-qPCR of whole FLOs on day 15, 24, and 30, hiPSCs and primary human bone marrow aspirate cells (BMACs, healthy donor) for embryonic (embr; *HBZ, HBE1*), fetal (*HBG1, HBG2*) and adult (*HBA, HBB*) hemoglobin genes, normalized *via* dCq to *RSP18*. Each dataset represents n=2-4 separate cultures for FLOs and n=2 technical replicates for BMACs.

**Supplementary Figure S2**

**A**: Representative brightfield images of Granulocyte, erythroid, macrophage and megakaryocyte colony forming units (CFU-GEMM) after 2 weeks culture in MethoCult SF H4436 (STEMCELL Technologies). Scale bar 500 µm. **B:** B-cell-like immunophenotyping per flow cytometry on unfractionated FLOs and umbilical cord blood (UCB) cells after 26 days of co-cultures on MS-5 cells, as well as primary adult spleen cells (*ex vivo*, not-cultured). Representative plots are shown of n=2 separate cultures. **C:** UMAP of Seurat-guided clustering of integrated FLO + GM/34 scRNA-seq datasets (days 10, 13, 16, 20). **D:** Dot plot with cluster-specific gene expression enrichment on nascent HSC-Scorecards per Calvanese et al. (Calvanese et al., 2022). **E**: Nascent HSC gene signature expression (Calvanese et al., 2022) of computationally isolated CD34 cells of FLO + GM/34 scRNA-seq datasets on days 10, 13, 16 and 20; embryonic fetal liver (FL) and yolk sac (YS) (Wang et al., 2021); and human colon organoid (HCO) hematopoietic cells (Múnera et al., 2023). **F:** UMAP feature plots of FLOs + GM/34 scRNA-seq datasets on days 13 and 16 of HPC and endothelial-to-hematopoietic transition (EHT) signature genes.

**Supplementary Figure S3**

Proximity of CD34+ CXCR4+ cells to HepPar1+ CXCL12+ assessed *via* **A**: Number of neighbors and **B:** Closest distance of CD34+ CXCR4+ cells within 100 µm radius of HepPar1 cells with (+/+) and without (+/−) CXCL12 co-localization based on mIF images. n = 8 separate organoid cultures shown as paired data points. Wilcoxon matched-pairs signed rank test (for A) and a paired two-tailed t-test (for B) was used to test for significance. p-values are shown. **C**: Percentage of CXCL12+ area within HepPar1+ cells assessed *via* mIF images on days 30 and 42. n=2 separate organoids shown as individual data points and mean ± SD. An unpaired two-tailed t-test was used to test for significance. p-values are shown. **D**: Quantification of CD45+ cells by flow cytometry of FLOs, and FLOs treated with CXCR4 antagonist AMD3100 (AMD) from day 26 for 2 weeks, relative to untreated FLOs. n = 2 separate organoid cultures shown as individual data points and mean ± SD. An unpaired two-tailed t-test was used to test for significance. p-values are shown. **E**: Granulocyte-macrophage colony forming units (CFU-GM) after 2 weeks culture in MethoCult SF H4436 of FLOs on days 22-27. Cells were plated either stromal cell depleted (CD34 enriched, CD34en), or unfractionated (whole). The colony numbers were normalized to 10000 plated CD34+ cells assessed *via* flow cytometry. n = 4 separate organoid cultures shown as individual data points min to max. An unpaired one-tailed t-test was used to test for significance. p-values are shown. **F**: Protein concentration of M-CSF on day 24 in FLO-, FLO + GM/34- and hiPSC-conditioned medium assessed *via* ELISA. Results are normalized to 1 mg total protein per 48 hours. n = 2-4 separate cultures shown as individual data points min to max. One-way ANOVA with Tukey multiple comparison test was used to test for significance. Corrected p-values are shown. **G**: Flow cytometry analysis of FLOs treated with and without CSF-1R-inhibitor BLZ945 (BLZ) and PBMCs. Gated for viable, single cells. Representative plots of n = 2 separate organoid cultures.

**Supplementary Figure S4**

**A**: UMAP demonstrating cell clusters (RNA_snn_res.0.8) of FLO + GM/34 scRNA-seq dataset on day 20. Top 5 expressed genes relative to all clusters shown for each cluster (Supplementary Table 4) **B**: Heatmap of day 20 cluster gene expression of gene ontology biological process (GOBP) genes for positive regulation of hematopoietic progenitor cell differentiation. **C**: UMAP of FLO + GM/34 scRNA-seq dataset on day 20 showing co-expression of indicated markers. lin =lineage. **D**: mIF image of day 42 FLO for HepPar1 (red), ligand IGF2 (yellow) and its receptor IGF2R (cyan). Representative image of n=8 separate organoid cultures. Scale bar 50 µm. **E**: IHC for stromal-hematopoietic niche per Nestin-staining on FLO + GM/34. Scale bar 40 µm.

**Supplementary Figure S5**

**A**: CytoTRACE rooted lineage trajectory analysis (Gulati et al., 2020) illustrates predicted root cells (red) as putative stem and progenitor cells and differentiation trajectories as yellow, green, and blue. **B**: UMAP of integrated FLO GM/34 scRNA-seq datasets (days 10, 13, 16, 20) with color-coded annotation to Human Fetal Liver atlas(Popescu et al., 2019), 2019 to illustrate multiple cell states found in human fetal liver; FOXA2+ cell population was after annotation defined as endoderm; HSC/MPP cluster was defined as HSPC. **C**: UMAP of FLO + GM/34 scRNA-seq dataset on day 16 annotated per two independent human developmental atlases, left fetal liver datssets (Popescu et al., 2019) compared to Fetal Liver (FL, center) and Yolk Sac (YS, right) (Wang et al., 2021). **D**: UMAP of FLO + GM/34 scRNA-seq dataset on day 16 integrated with FL and YS datasets(Wang et al., 2021). **E**: Clustering of integrated datasets from day 16 FLO + GM/34, FL and YS (left) with percentage of contribution to respective clusters (right). **F**: IHC of FLO + G/GM/34 (added G-CSF to FLO + GM/34 from day 20) on day 30 and fetal liver (pcw 39) of myeloid markers CD68, CD15, and neutrophil elastase (NE), as well es stromal markers CD31 and aSMA. Representative images shown of n=6 separate organoid cultures. Scale bar 100 µm. **G**: PCA plot of bioinformatically isolated CD45+ cells of FLO + GM/34 scRNA-seq dataset on day 16 integrated with datasets from human fetal macrophage development (Bian et al., 2020).

**Supplementary Figure S6**

**A**: Expression heatmap of bulk RNAseq data and unbiased, hierarchical sample clustering for a monocyte-related gene panel. Samples include primary cells, FLO + GM/34-derived myeloid subsets (magenta), and other iPSC-derived immune cells. The primary cells include LPS and IL4/IL13-polarized PBMC-derived macrophages, primary PBMC-derived neutrophil granulocytes (Prim Neutro Granulo), primary PBMC-derived monocytes (Class=classical, Int=intermediate, and NC=nonclassical subsets, monocyte data derived from Monaco et al.(Monaco et al., 2019) and integrated into our dataset), CD68+ adult liver macrophages, iPSC-derived neutrophil granulocytes and -monocytes differentiated as per Trump et al. (Trump et al., 2019) (iPSC-Neutro Granulo and -Mono, respectively). Every sample represents a separate culture. **B-D**: mIF images of the mouse kidney 2 weeks (2w) after transplantation of FLOs into the mouse kidney capsule (KC). Scale bar 500 µm (B) and 50 µm (C, D). **E**: mIF image of the mouse kidney 2 weeks (KC1, left) and 6 weeks (KC4, right) after transplantation of FLOs (using *RUNX1* murine +23 donor dsRed enhancer reporter ESC line) detecting NM95+ CD34+ hCD45+ cells (top) and dsRed signal using RFP antibody (bottom). Scale bar 20 µm. **F**: Quantification of mIF images for CD34 (left) and hCD45 (right) relative to NM95+ cells of the mouse kidney from untransplanted (NC1), 2 weeks (KC1) and 6 weeks (KC4) transplantation into the KC. n=2 technical replicates per mouse shown as individual data points and mean ± SD. **G**: Quantification of mCD45-hCD45+ cells (left) and mCD45-hCD45+ hCD34+ cells (right) in mouse KC 2 weeks after FLO transplantation assessed *via* flow cytometry. Frequencies of viable single cells shown. Each color represents one KC transplanted mouse (KC2 pink, and KC3 brown). n=2 transplanted mice shown as individual data points and mean. **H**: mIF image of KC4, 6 weeks (6w) after FLO-transplantation. Arrows highlighting capillary-like structures. Scale bar 50 µm. **I**: Flow cytometry (FC) of mouse liver 6 weeks after transplantation into the KC to distinguish mouse (m) and human (h) CD45+ cells. Gated for viable single cells (left) and mCD45-hCD45+ (right).

**Supplementary Table 1:** PSC-lines used in this study

**Supplementary Table 2:** List of Oligonucleotides for RT-qPCR

**Supplementary Table 3:** Top expressed genes of each cluster relative to other clusters of day 13 FLO+GM/34.

**Supplementary Table 4:** Top expressed gene list of each cluster relative to other clusters of day 16 FLO+GM/34.

**Supplementary Video 1:** Live cell imaging of FLOs made from iPSC72_3-AAVS1-CAG-GFP-line, releasing proliferative blood cells.

**Supplementary Video 2:** 3D-light sheet microscopy of FLO on day 36 after whole-mount staining with fluorescent CD34-antibody (magenta), CD31- (yellow), and ECAD- (green) nanobody.

## EXPERIMENTAL MODELS AND SUBJECT DETAILS

### Animals

Immune-compromised NOD-SCID IL-2Rgnull (NSG) mice (between 8-12 weeks old) for transplantation studies were obtained from the Comprehensive Mouse and Cancer Core Facility in Cincinnati, Ohio. All experimental procedures were conducted with approval from the Institutional Animal Care and Use Committee (IACUC) at CCHMC.

### Cell lines

#### Human pluripotent stem cell (hPSC) lines and maintenance

Human PSC lines listed in Supplementary Table 1 were maintained as described previously(Ouchi et al., 2019; Shinozawa et al., 2021). Briefly, undifferentiated hPSCs were maintained on feeder-free conditions in mTeSR1^TM^ (STEMCELL Technologies) on 6-well plates (BD Falcon) coated with Geltrex (Thermo Fisher, 1:120 in KO DMEM, 30 min at 37 °C), Vitronectin (Thermo Fisher, 1:100 in PBS, 1 h at RT), or Laminin (iMatrix-511, 1% in PBS, 2 h at 37 °C). Frozen hPSCs were thawed in mTeSR1^TM^ with the addition of Y27632 (STEMCELL Technologies, 10 µM). hPSCs were passaged every 3-5 days using EDTA (0.5 mM in PBS, 3-5 min at room temperature) for dissociation. Cells were seeded as clumps in a 1:10 to 1:20 ratio onto new coated plates as described above. For single-cell seeding, cells were dissociated using Accutase (Thermo Fisher) or TrypleE Express (Thermo Fisher) for 5 min at 37 °C and seeded onto coated plates in mTeSR1^TM^ with Y27632.

Cells were maintained at 37°C in 5% CO2 with 95% air, and the medium was replaced daily. To ascertain genomic integrity, standard metaphase spreads and G-banded karyotypes were determined by CCHMC Cytogenetics Laboratory and BIH Stem Cell Core of the used PSC-lines (Sup. Table 1).

#### Generation of human PSC reporter line

For the generation of H9 (WA09, WiCell) *RUNX1* murine +23enh dsRed reporter line zinc finger nucleases were used as detailed in Fig. S1F to introduce a DNA double-stranded break into the first intron of the *PPP1R12C (AAVS1)* locus. The donor matrix containing two homologous arms flanking a splice acceptor, 2A cleavage side, puromycin resistance gene followed by the murine +23 enhancer and dsRed fluorescence reporter gene driven by Hsp68 promoter sequence was assembled using In-Fusion® HD Cloning Kit (Clontech Laboratories Inc.). The Hsp68 promoter dsRed sequence and downstream murine +23 enhancer were integrated into the opposite orientation of the *PPP1R12C* gene locus, which resulted in a stronger fluorescence signal compared to the integration of the reporter in orientation with *PPP1R12C*. The 531-bp mouse RUNX1/AML1 +23 enhancer was chosen due to its high gene homology with homo sapiens and previously published work that identified this enhancer element being able to drive hematopoietic cell emergence in a *Runx1*-specific fashion(Bee et al., 2009; Nottingham et al., 2007). Donor DNA and bicistronic AAVS1 zinc finger nucleases were delivered into H9 using nucleofection as per the manufacturer’s instructions (Lonza Bioscience). Electroporated single cells were seeded into Corning^®^ Matrigel^®^-coated cell culture plates and cultured in mTeSR1™ (STEMCELL Technologies Inc.) supplemented with 10 μM Y-27632 (Tocris) for 1-2 days before adding puromycin (InvivoGen) to select for correctly targeted cells. Puromycin-resistant colonies were transferred into individual wells to generate clonal populations. Successful homologous recombination and donor cassette integration were confirmed by targeted PCR followed by sequencing. Verified clones were expanded and biobanked. Genomic stability was confirmed by karyotyping.

### Primary human specimens

*Umbilical cord blood (UCB)* units were collected by the Translational Trials Development Support Laboratory of Cincinnati Children’s Hospital Research Foundation and distributed according to an approved IRB protocol, and selected for CD34^+^ cells with the EasySep human CD34^+^ magnetic isolation kit (STEMCELL Technologies).

*Bone marrow aspirate cells (BMACs) and CD34-selected cells from mobilized peripheral blood stem cells (mPBSCs)* were obtained from allogenic hematopoietic stem cell (HSC) transplantations at Charité Universitätsmedizin Berlin.

*Peripheral blood mononuclear cells (PBMCs) and granulocytes* were obtained from peripheral blood of a healthy, male, adult donor under consent, and isolated using Ficoll-Paque Premium (GE Healthcare Life Sciences).

For PBMC-derived macrophage derivation, we adapted a published protocol (Schwager et al., 2022). To generate inflammatory macrophages, PBMCs were incubated for 6 days with GM-CSF (25 ng/ml, STEMCELL Technologies) and then polarized with LPS (100 ng/ml, Sigma) for 24 hours. To generate anti-inflammatory macrophages, PBMCs were incubated with M-CSF (25 ng/ml, STEMCELL Technologies) and then polarized with IL4 (40 ng/ml, R&D Systems) and IL13 (40 ng/ml, Peprotech) for 24 hours.

*Human fetal liver specimens* archived, autopsy-derived, anonymous FFPE-fixed tissue sections were provided by the CCHMC Pathology core (16, 18, and 39 post conception weeks).

*Postnatal liver specimens* were obtained from pediatric liver explant or biopsy at Charité Universitätsmedizin Berlin under parental informed consent.

*After informed consent, primary adult hepatocytes (AHs)* were obtained from adult surgical liver explants at Charité Universitätsmedizin Berlin (Department of Surgery).

*IRB-exempt, brain-dead donor spleen samples* were received from the laboratory of Appakalai N. Balamurugan under research consent. For splenocyte extraction, fresh human spleen specimens were cut into small pieces approximately 2cm^3^ and placed on a 10 cm petri dish on ice. 1000 µl of cold PBS (Sigma) was added to the plate. A 50 ml tube with about 10 ml cold PBS was added to the ice and a second, empty 50 ml tube was also added to the ice. Spleen pieces were ground between two microscope slides with frosted sides against each other to provide friction. The spleen was rinsed off the microscope slide with a 1000 µl pipette and the resulting liquid was passed through a 100 µm filter into the 50 ml tube with 10 ml PBS. The process was repeated with multiple pieces as necessary. After all pieces had been processed, the product was then passed through a 40 µl filter into the empty 50 ml tube. The tube was centrifuged at 300g for 5 minutes at 4°C. PBS was aspirated and cells were either used immediately in flow cytometry experiments or frozen at −80*C in 90% FBS (HI, LifeTechnologies) with 10% DMSO (Sigma).

## METHOD DETAILS

### Differentiation of hPSCs into FLOs

hPSCs were cultured for at least 2 passages after thawing in mTeSR1^TM^ (STEMCELL Technologies) before differentiation. After reaching 80% confluency, hPSCs were seeded as single cells (2-4×10^5^ cells/ml, see hPSC maintenance) on coated 12-well plates (BD Falcon). Cells were constantly maintained at 37°C in 5% CO2 with 95% air.

After 24 hours, Y27632 was removed. For co-induction of endoderm and hemogenic mesoderm, medium was changed to RPMI 1640 medium (ThermoFisher) containing 100 ng/ml Activin A (Peprotech) and 50 ng/ml BMP4 (R&D Systems) at day 0; 100 ng/ml Activin A and 0.2% FBS (Cytiva) at day 1, and 100 ng/ml Activin A and 2% FBS on day 2. From day 3 on, cells were maintained in basal media consisting of advanced DMEM/F12 (Gibco) supplemented with 2% B27 (Gibco), 1% N2 (Gibco), 1% GlutaMAX (Gibco), 1% HEPES (Gibco), and 1% Pen-Strep (Gibco). During the posterior foregut stage (day 3-5), basal media was supplemented with 500 ng/ml FGF4 (Peprotech) and 3 µM CHIR99021 (Hellobio) and the media was changed every day.

On day 6, cells (> 90% viability) were washed in PBS and collected in clusters by pipetting 5-7 times in advanced DMEM/F12, followed by centrifugation (3 min, 300 g) and media aspiration. For each well of a 12-well plate, 500 µl Matrigel (Corning, Standard formulation, diluted to 8.8 mg/ml protein in advanced DMEM/F12) was added. Cells were resuspended by gentle pipetting, not more than 10 times to maintain cell clusters. All steps including Matrigel were performed on ice. Then, 50 or 75 µl Matrigel-cell suspension drops were placed in the center of each well of a 24-well plate (VWR), and air bubbles were avoided. After incubation at RT for 3 min, the plate with Matrigel domes was flipped upside-down to avoid sinking of the cell clusters to the bottom and incubated at 37 °C for additional 15 min for Matrigel solidification. Next, plates were flipped back, and 600 µl/dome basal medium supplemented with 20 ng/ml BMP4 and 10 ng/ml FGF2 was added and incubated for 4 days. From day 10, only basal medium without growth factors was added to ensure self-organization and changed every 3 days, and from day 22 every 2 days.

### Modulations in FLO differentiation

To enhance myeloid differentiation, 50 ng/ml IL-3 (STEMCELL Technologies), 50 ng/ml IL-34 (Biolegend), 50 ng/ml GM-CSF (STEMCELL Technologies), and 35 nM UM171 (Apexbio Technology) or 500 nM UM729 (STEMCELL Technologies) was added from day 10, and referred to as “GM/34”. For experiments involving granulocyte maturation, 100 ng/ml G-CSF (STEMCELL Technologies) was additionally supplemented from day 20.

To assess functionality of FLO-derived CXCL12, CXCR4 antagonist AMD3100 (Santa Cruz, 100 µg/ml) was added to the FLO basal medium from day 26.

To assess functionality of FLO-derived CSF, CSF-1R inhibitor BLZ945 (Hycultec, 500 nM) was added to the basal medium from day 10.

For transplantation experiments, we applied 5 ng/ml FGF2 (R&D Systems), 20 ng/ml EGF (R&D Systems), 10 ng/ml VEGF (Life Technologies), 3 µM CHIR (R&D Systems), 10 mM A83-01 (Tocris) and 50 µg/ml ascorbic acid from day 7-10, and 10 ng/ml FGF2 (R&D Systems) and 20 ng/ml BMP4 (R&D Systems) from day 10-14.

### FLO transplantation into NSG mice

NSG mice were kept on standard chow. Food and water were provided ad libitum before and after surgeries. Transplantations were conducted between 8 to 12 weeks old mice. Day 21 FLOs were removed from Matrigel, washed with cold phosphate-buffered saline (DPBS; Gibco), and transplanted under the kidney capsule (KC) or injected intraperitoneally (IP). Mice were anesthetized using 2% inhaled isoflurane (Butler Schein), and the left side was sterilized with isopropyl alcohol and povidone-iodine. A small incision was made in the left posterior subcostal region to expose the kidney. A subcapsular pocket was created, and the organoid was placed inside. The kidney was then returned to the peritoneal cavity. The skin was sutured in two layers, and the mice received a subcutaneous injection of Buprenex (0.05 mg/kg; Midwest Veterinary Supply). Transplanted mice were euthanized and dissected 2 and 6 weeks after transplantation. Transplanted kidneys, liver, peritoneal cells, spleen, and bone marrow cells were collected for FFPE processing, or made into single-cell suspension for flow cytometry for markers hCD34, hCD45 and mCD45.

### Embryoid body-based directed differentiation from hPSCs into HSPCs (EB-HSPCs)

For Colony Forming Unit (CFU) assays we here used an embryoid body (EB)-based differentiation protocol adapted from a published protocol (Atkins et al., 2022)a. All differentiation steps were performed in StemPro-34 media (Gibco) supplemented with 2 mM GlutaMAX (Gibco), 1% Pen-Strep (Gibco), 50 µg/ml ascorbic acid (Sigma), 150 μg/ml transferrin (Roche), 50 μg/ml monothioglycerol (Sigma). On day 0, hPSCs were singularized at 80% confluency using TrypleE Express (Thermo Fisher), and 750K cells/ml were seeded in media containing 10 µM Y27632 and 1 ng/ml BMP4 (R&D Systems) on 24-well AggreWell400 (STEMCELL Technologies) pre-treated with Anti-Adherence Rinsing Solution (STEMCELL Technologies). After incubation for 18-24 hours at 37 °C, EBs from each AggreWell were transferred to an Anti-Adherence Rinsing Solution treated well of a 12-well plate in media with 10 ng/ml BMP4 (R&D Systems), 5 ng/ml FGF2 (R&D Systems), and 6 ng/ml Activin A (Peprotech). On day 4, media was replaced with 5 ng/ml FGF2 (R&D Systems), 15 ng/ml VEGF (Peprotech), 10 ng/ml IL-6 (Peprotech), and 10 ng/ml IL-11 (Peprotech). From day 6 onward, cultures were maintained in media containing 5 ng/ml FGF2 (R&D Systems), 15 ng/ml VEGF (Peprotech), 10 ng/ml IL-6 (Peprotech), and 10 ng/ml IL11 (Peprotech), 100 ng/ml SCF (Peprotech), 50 ng/ml IGF1 (Peprotech), and 4 U/ml EPO (Peprotech) with media changes every other day. On days 9-10, the cells were harvested for CFU assays.

For T-cell assay, differentiation of hPSCs to HSPCs was performed using the StemDiff T Cell Kit (STEMCELL Technologies) following the manufacturer’s protocol with the following modifications. In short, hPSCs were harvested as single cells and seeded at 3.6×10^5^ cells/well on a 24-well AggreWell400-plate (STEMCELL Technologies) in 2 ml EB Medium A supplemented with 10 µM ROCK inhibitor Y-27632 (STEMCELL Technologies) and incubated at 37°C, 5% CO2 to generate EBs. Half of the medium was exchanged with fresh EB Medium A on day 2, and with EB Medium B on day 3 of culture. On day 5, EBs were collected from the AggreWell plate and pooled in a 37 µm reversible strainer (STEMCELL Technologies). EBs were washed off with fresh EB Medium B and seeded in a non-tissue culture-treated 24-well plate with 1 ml medium per well. 1 ml EB Medium B was added on day 7 of culture, and another half-medium exchange was performed on day 10. On day 12, EBs were harvested for immunophenotyping and T-cell assay.

### Harvesting and enrichment of CD34 cells

FLOs and EB-HSPCs were pooled and dissociated using 2.5 mg/ml Collagenase IV (Worthington), 1 mg/ml DNase I (Merck) in HBSS (Gibco), followed by straining (40 µm) to obtain a single-cell suspension. CD34 cells were enriched using the CD34 Microbead kit (Miltenyi) according to the manufacturer’s protocol. To increase purity in FLOs, the enrichment step was repeated.

### Colony Forming Units (CFU) assay

Unfractionated (1×10^4-5×10^5 cells) or CD34-enriched (1×10^3-1×10^4 cells) single-cell suspensions were plated onto 35mm culture dishes in MethoCult SF H4436 (STEMCELL Technologies), and incubated at 37°C, in 5% CO2 with ≥ 95% humidity. After 14 days, brightfield images were taken with the Keyence BZ-X800. Colonies were counted and normalized to 10k plated CD34+ cells assessed *via* flow cytometry.

### Lymphocyte differentiation assays

CD34-enriched cells from FLOs, FLOs + GM/34, EB-HSPCs, and primary mPBSCs from a healthy donor (female, 30 years) were differentiated using the StemDiff T Cell Kit (STEMCELL Technologies) with minor modifications to the manufacturer’s protocol. Between 0.5 – 1×10^5^ cells were seeded onto one well of a non-tissue culture-treated 6-well plate coated with lymphoid differentiation coating solution (LDCS) and cultured in 5 ml StemSpan Lymphoid Progenitor Expansion Medium (LPEM) per well. A half-medium exchange of LPEM was performed every 3-4 days and non-adherent cells were transferred to a freshly LDCS-coated well at day 6 of culture. After 14 days of culture, differentiated non-adherent cells were harvested and re-seeded at 2×10^5^ cells/well on LDCS-coated non-tissue culture-treated 24-well plates and cultured with 1 ml StemSpan T Cell Progenitor Maturation Medium (TPMM). Half-medium exchange with fresh TPMM was done every 3-4 days until flow cytometry analysis on differentiation day 28 for T-cell markers CD5, CD7, CD8α, CD8β, CD3, TCRαβ and CD4.

For B-cell induction, we adjusted the protocol of Richardson et al., 2021(Richardson et al., 2021) by adding either umbilical cord blood (UCB) or CD34+ sorted FLO-cells to MS-5 co-culture, followed by flow cytometry on day 26 for CD45, CD20, CD43 and CD27.

### Migration Assay of CD34+ cells

To evaluate the functionality of CXCL12 secreted by FLOs, a migration assay was performed using CD34+ cells derived from mPBSCs. Conditioned medium was collected from FLO cultures on day 22 after a 3-day incubation period, as well as from hiPSCs, which were used as a control. All media were stored at −80°C until further use.

For the migration assay, conditioned media from FLOs and hiPSCs were added to 24-well plates. Unconditioned media were used as baseline controls. Positive control included media supplemented with SDF-1 (R&D Systems, 100 ng/ml), while CXCR4 inhibitor AMD3100 (Santa Cruz, 0.1 mg/ml) was added to FLO-conditioned media to assess CXCL12/CXCR4-dependent migration. Media were added to the wells, and 6.5 mm Transwell inserts with a 5.0 µm pore size (Corning) were placed in each well. CD34+ cells (50,000 cells per 100 µl) were then added to the upper chamber of the Transwell inserts. CD34+ cells were thawed 30 minutes before the assay and kept in advanced DMEM at 37°C and 5% CO2. After a 4.5-hour incubation period, the Transwell inserts were removed, and the migrated cells were counted using flow cytometry by recording events per two minutes. Blank samples were subtracted from the conditioned media samples during data analysis.

### Depletion assay

For cell depletion assays, we performed FLO induction as described before. On day 6, we singularized the cells for FACS-sorting using Accutase (Sigma). We individually stained cells for KDR, EPCAM or NCAM according to manufactureŕs instructions and included DAPI (Invitrogen) as viability stain for all samples. We then sorted respective positive and negative populations pre-gated for single & viable cells using BD AriaFusion with an 85 µm nozzle (BIH Flow & Mass Cytometry Core Facility). Subsequently, we either seeded cells in Matrigel domes as isolated negative populations or pooled together with respective positive populations according to the initial ratio. Matrigel volume was normalized to cell number assuring all cells had the same seeding density (100k cells per 50 µl Matrigel). Then, cells underwent the standard FLO protocol.

### ELISAs for CXCL12, EPO, M-CSF

We measured secreted protein levels in FLOs and respective control cultures for CXCL12, EPO and M-CSF using commercial Human CXCL12/SDF-1 alpha Quantikine ELISA (R&D Systems), LEGEND MAX™ Human Erythropoietin (EPO) ELISA (BioLegend) and Human M-CSF (CSF-1) ELISA (Invitrogen) kits, respectively, according to manufactureŕs instructions. We collected supernatants from cultures after 48 h of culture. These we centrifuged and then froze and stored in −80°C until performing the assay. We used undiluted supernatants for the assay. We normalized ELISA results to the total protein content assessed using the Pierce™ BCA Protein Assay Kit (Thermo Scientific). For this we measured serial dilutions of the total protein content of respective cultures.

### Cytochrome activity (CYP3A4) assay

To measure cytochrome activity, the Promega #V8901 (luciferin PFBE) kit was used as CYP3A4 activity assay according to manufacturer’s instructions. Cell viability or count was assessed for normalization. In one experiment, CYP3A4 activity in day 30-FLOs was assessed relative to primary human hepatocytes cultured for 24h on collagen-coated plates (HepaCur, Cryopreserved) In another experiment, CYP3A4 activity in iHeps (conventional PSC-derived differentiation protocol into hepatocyte-like (Kajiwara et al., 2012) was assessed relative to primary human hepatocytes from liver explant (frozen).

### Phagocytosis assay

To assess phagocytosis, the respective myeloid immune cells were isolated by FACS-based cell sorting via Sony/SH800S using a 100 µm nozzle with the respective antibodies. Isolated cell subsets were incubated for 6 hours on Fibronectin-coated (ThermoFisher) cell culture plates, followed by incubation with pHrodo™ Deep Red E. coli BioParticles™ Conjugate for Phagocytosis (ThermoFisher) according to the manufacturer’s instructions by adding particle stock solution after vortexing directly to the media. The phagocytic activity was quantified by fluorescent microscopy-based imaging after NucBlue™ staining (ThermoFisher).

### Reactive oxygen species assay

To assess ROS generation, Granulocytes cultured on Fibronectin-coated plate (ThermoFisher) were treated with 100 nM of Phorbol 12-myristate 13-acetate (PMA; Tocris) before performing the OxiSelect™ In Vitro ROS/RNS Assay Kit (CellBio Labs).

### Granulocyte migration assay

To assess granulocyte migration, neutrophils were collected by FACS from FLOs + GM/34 or by FICOLL cell prep (see above) and migration was assessed towards a fMLP-containing media (33 mM, Sigma) using the CytoSelect 96-Well Cell Migration Assay (Cell Biolabs) after 60 minutes at 37°C for in 5% CO_2_ cell culture incubator. Cells that had migrated were counted using a conventional hemocytometer.

### Inflammatory cytokine secretion assay

For the inflammatory cytokine profiling using electrochemiluminescence assay, the experiment was carried out by collecting the supernatant from organoids treated with various conditions. 24 hours-Lipopolysaccharide (LPS)-challenged conditions were included (Sigma-Aldrich, St. Louis, Missouri, United States). The sample was used fresh or stored at −80 C for future use. Quantification of 6 proinflammatory cytokines (IFN-γ, IL-1β, IL-6, IL-8, IL-8 (HA), and TNF-α) was performed using V-Plex Proinflammatory Panel 1 Human Kits and V-Plex Chemokine Panel 1 Human Kit (Meso Scale Diagnostics). Samples were analyzed in duplicates or triplicates and assays were performed per manufacturer protocol and read and analyzed on a QuickPlex SQ 120MM (Meso Scale Diagnostics). Briefly, all reagents, and samples were thawed on ice; the calibrator was reconstituted and incubated for 15-30 min at room temperature. Sample, standard and calibrators were diluted as required. The plates were then washed with 1X PBST (PBS/ 0.05 % Tween-20) three times. The calibrators and samples were added to separate wells of both plates and incubated for 2 hours with shaking (500 – 700 RPM) at room temperature. The plates were again washed with 1X PBST three times and the detection antibodies were added to e ach well and incubated for 2 hours with shaking (500 – 700 RPM) at room temperature. The plates were washed one final time with 1X PBST three times. The read buffer was then added, and the plate was read immediately on a QuickPlex SQ 120MM and analyzed with DISCOVERY WORKBENCH v4.0.013 (Meso Scale Diagnostics, Rockville, MD, USA).

### Flow cytometry

For surface marker flow cytometry, we stained cells in PBS containing 0.2% BSA, 2 mM EDTA and the antibody dilutions according to manufactureŕs instructions. For intra-cellular marker, we used the Transcription Factor Staining Buffer Set (Miltenyi) according to manufactureŕs instructions. Unspecific binding sites were blocked for spectral flow cytometry by incubation with 1 mg/ml Beriglobin and for 3-laser based cytometry using BSA. We analyzed cells using the Aurora spectral cytometer (Cytek Biosciences) or BD FACSCanto™ II (3-laser). For flow cytometry analysis of transplantation experiments, 10^6^-10^7^ cells were suspended and stained in 100 μl staining buffer (2 mM EDTA and 5% FBS in PBS) and analyzed using FACS Melody. Subsequent analyses were performed using FlowJo™.

### Giemsa Staining

For Giemsa, 7000 GYPA+ sorted cells or cells from CFU assays collected from methylcellulose by washing with PBS, were cytospun onto slides (500 rpm for 10 min). Slides were air-dried and stained with a Wright-Giemsa staining kit (ThermoFisher), according to the manufacturer’s instructions followed by examination by light microscopy. For the manual differential cell count of sorted FLO + GM/34 cell subsets a hemopathologist, who was blinded for the different conditions, performed a manual count.

### Immunohistochemistry FFPE samples

For Immunohistochemistry (IHC), samples in Matrigel drops were collected, fixed in 10% formalin, and embedded in paraffin. 5 µm sections were subjected to H&E and immunohistochemical staining.

### Multiplexed immunofluorescence (mIF)

mIF was performed on 5-μm thick formalin-fixed paraffin-embedded organoid sections as previously described(A et al., 2020; Guillot et al., 2020). Briefly, antigen retrieval was performed in heated citrate buffer (pH=6.0). Antibodies were eluted after each cycle of staining using a 2-mercaptoethanol/SDS (2-ME/SDS) buffer. Primary and secondary antibodies used in this study are listed in the Key Resources Table. Images were acquired on a Zeiss AxioObserver 7 and processed in FIJI as previously reported(Guillot et al., 2023).

### Whole-mount organoid immunofluorescence staining for light sheet microscopy

Organoids fixed in 4% PFA were permeabilized (0.5% Triton X-100 in PBS), then blocked with PermBlock solution (1% BSA, 0.5% Tween 20 in PBS). Whole-mount immunofluorescence staining was performed using indicated primary antibodies and Alexa dye-coupled secondary antibodies diluted in PermBlock solution. After each staining step, samples were washed three times in PBS-T.

### Optical clearing of whole-mount-stained organoids

The immunofluorescence-stained organoids were embedded in 1% low-melting-point agarose (Thermo Fisher Scientific), then dehydrated through a series of increasing methanol concentrations (50%, 70%, 95%, >99.0%, >99.0% [v/v] methanol, each step for 30 minutes). The samples were subsequently optically cleared twice in a BABB solution (benzyl alcohol:benzyl benzoate solution, ration 1:2). The optically cleared organoids were stored in BABB until imaging.

### Light sheet microscopy and 3D reconstruction

Immunofluorescence-stained and optically cleared organoids were optically sectioned using a Lightsheet 7 microscope (Zeiss). Images stacks were acquired with a step size of 1.5 µm at various magnifications. The digital three-dimensional reconstruction of light sheet image stacks and respective rendering was conducted using the IMARIS Microscopy Image Analysis Software (Oxford Instruments).

### Image analysis

Binary masks were generated using the trainable classification software Ilastik (v1.3.3). Neighbor analysis was performed on CellProfiler.

### RNA isolation and RT–qPCR

To isolate RNA, samples were homogenized using 1 ml Trizol (Thermo Fisher), followed by addition of 0.2 ml chloroform and vortexing. After centrifugation for 10 min, 12,000 g at 4°C, the aqueous phase was transferred to a new tube and mixed with 70% ethanol 1:1. Subsequent column purification steps were conducted with RNeasy mini kit (Qiagen) following the manufacturer’s instructions. The concentration of the isolated RNA was assessed using Nanodrop. cDNA was generated using High-Capacity cDNA Reverse Transcription Kit (Thermo), cDNA Synthesis Kit (biotechrabbit^TM^), or High-Capacity RNA-to-cDNA™ Kit (Applied Biosystems) according to the manufactureŕs instructions. RT-qPCR was performed using the PowerUp SYBR Green Master Mix (Applied Biosystems) or TaqMan gene expression master mix (Applied Biosystems) and probes from the Universal Probe Library (Roche) on a QuantStudio 3 Real-Time PCR System (Thermo Fisher) according to manufacturer’s instructions. The primers were synthesized by Eurofins Genomics. Primer sequences are listed in Supplementary Table 2. Gene expression changes were analyzed using the ddCT method or dQC method as indicated.

### Bulk RNA processing

The FASTQ files from paired-end sequencing were initially processed (including trimming, filtering low-quality reads, and removing adaptors) using the command line tool fastp (Chen et al., 2018). Quantification is performed via pseudo alignment using the R package Kallisto against the human reference transcriptome hg38(Bray et al., 2016). Gene counts are then estimated from the resulting transcript abundances using the R package tximport(Soneson et al., 2015), and samples are combined into a final counts matrix. The function mnnCorrect from the R package Batchelor was adapted to correct the counts for batch effects using a mutual nearest neighbors method (Haghverdi et al., 2018). Differentially expressed genes were then identified between populations of interest using R package DESeq2(Love et al., 2014) and visualized using volcano plots. The corrected counts matrix is log-transformed for principal component analysis, and principal components are calculated and visualized. The top and bottom 100 genes from the first four principal components are subsets from the normalized counts and used for generating the expression heatmap.

### Single-cell RNA-seq sample handling and processing

FLOs + GM/34 on day 10, 13, 16 and 20 were singularized with Trypsin, cryopreserved and sent to GENEWIZ/Azenta Life Sciences (New Jersey, USA). There, the manufacturer’s instructions for scRNA-sequencing were subsequently followed utilizing 10x Genomics Chromium; cells and barcoded gel beads were separated in partitioning oil droplets. Cell- and transcript-specific barcodes were added during reverse transcription. The Illumina platform was used to pool and sequence barcoded libraries. FASTQ files were first aligned against the human hg38 genome using Cellranger, outputting a counts matrix, estimated number of cells, number of reads and genes per cell, and flags for any quality issues detected (https://github.com/10XGenomics/cellranger). SoupX is used to further process the counts by correcting for ambient RNA (https://academic.oup.com/gigascience/article/9/12/giaa151/6049831?login=true), which is then read in and processed by the R package Seurat (https://www.cell.com/cell/fulltext/S0092-8674(19)30559-8) (Stuart et al., 2019). An initial filtering step retains genes expressed in a minimum of 3 cells and cells expressing a minimum of 100 genes. *NormalizeData* then performs a per-cell global scaling normalization to adjust and log-transform the expression values. Feature selection with *FindVariableFeatures* ranks genes by their variability across cells while controlling for their mean expression, and the top 2000 genes are used for downstream analysis. Counts are also scaled with *ScaleData* to have mean 0 and variance 1, and *RunPCA* performs principal component analysis, a form of linear dimensionality reduction. Using the first 20 principal components from the PCA, *FindNeighbors* and *FindClusters* generate a shared nearest neighbor graph and identify clusters of cells at a resolution of 0.8. To visualize the clusters, *RunUMAP* is run to generate the uniform manifold approximation projection, a non-linear dimensionality reduction of the data.

### Cell type annotation

To identify cell populations, present in the samples in an automated approach, the R package ELeFHAnt was used (https://www.biorxiv.org/content/10.1101/2021.09.07.459342v1.full) (Thorner et al., 2021). ELeFHAnt utilizes a supervised machine learning approach where two classifiers, random forest (RF) and support vector machine (SVM), form an ensemble that trains on annotated single-cell RNA-seq data to classify “test” datasets. For our purposes, a reference composed of fetal liver cells (https://www.nature.com/articles/s41586-019-1652-y)(Popescu et al., 2019) served as the training dataset (∼150k cells) and the samples as the test datasets. The *ClassifyCells* function down-sampled the reference to 500 cells per cell type and trained on the 2000 most variable shared features across all datasets (using the parameters *downsample_to* and *selectanchorfeatures).* Cell types were then predicted for each cell using the ensemble learning method.

### Integration of samples to understand similarity and differences across cell populations

The four time points were integrated into a single dataset with Seurat for joint analysis. The function *FindIntegrationAnchors* takes two datasets and performs dimensionality reduction using diagonalized canonical correlation analysis (CCA), followed by L2-normalization of the canonical correlation vectors, and finding mutual nearest neighbors (MNNs) in this low dimensional space. The function *IntegrateData* uses these neighbors or “anchors” to calculate a weighted correction vector that reflects the batch effect. The batch effect is removed by subtracting these correction vectors from the expression matrix. For more than two datasets, such as in our case, this repeats in a recursive pairwise fashion until all are corrected. The integrated dataset can then be clustered and visualized (using the functions *RunPCA, FindNeighbors, FindClusters* and *RunUMAP*). When integrating with outside datasets, this process was repeated, but datasets were re-normalized using *SCTransform* (a method based on Pearson residuals) in place of *NormalizeData.* All single cell objects were also updated to the Seurat v5 format.

The integrated dataset resolves conflicting annotations with the function *LabelHarmonization* from ELeFHAnt. Taking the individual annotations for each dataset produced by *ClassifyCells*, it finds a shared set of annotations to annotate the integrated data. The method differs in that the integrated data serves as both the training and test sets, but otherwise, the same ensemble learning method with the same down-sampling (1000 cells per cell type) and feature selection (2000 integration features). Following training and classification, the cell type with the highest cell votes is assigned to their respective integration cluster.

### scRNA projection onto reference cell atlas

To further confirm the identity of cells present in the four samples, the *MapQuery* function from Seurat was used. In a process similar to integration, pairs of cells are found between the samples and a multi-organ fetal atlas reference (https://www.science.org/doi/10.1126/science.aba7721) (Cao et al., 2020) by projecting the reference PCA onto each query. Using these anchors, a weighted vote classifier annotates the cells in each query with the best matching reference cell type. Finally, the predictions are visualized by projecting each query onto the reference UMAP.

### Inferring trajectory across cells from 4 timepoints

To elucidate the cell states and if exists a trajectory across these states, we deployed dyno package in R (https://github.com/dynverse/dyno). Dyno serves as an interface to 60 different trajectory inference methods for scRNA datasets. We used dyno’s guidelines Shiny app, based on benchmarking results of trajectory inference methods, to obtain the optimal set of methods, which can be applied on our dataset. We applied following three methods: 1) PAGA (https://github.com/theislab/paga) 2) PAGA Tree (https://github.com/dynverse/ti_paga_tree) and 3) Slingshot (https://bmcgenomics.biomedcentral.com/articles/10.1186/s12864-018-4772-0). For PAGA and PAGA Tree cells in day 10 timepoint were set as root cells while slingshot uses an unsupervised approach to find root and terminal states. PAGA and PAGA tree use a topology preserving map while slingshot applies a minimum spanning tree to partition the cells to derive root, migrating and terminal fates.

### CytoTRACE

We applied CytoTRACE (Gulati et al., 2020) to reconstruct the developmental trajectory across cells from Dadays0 to 20. CytoTRACE determines the development potential of cells, modeling the number of genes expressed per cell; as cells mature, the transcriptional diversity/number of genes expressed decreases. Gene counts across cells from each timepoint were inputted to the *iCytoTRACE* function, which leverages mutual nearest neighbor and Gaussian kernel normalization techniques to merge datasets and correct gene expression. Following correction, it calculates the development potential of cells based on available transcriptional diversity.

### Statistical Software

Statistical analyses were mainly performed using GraphPad Prism 9.0 (GraphPad Software, Inc., CA, USA), using appropriate tests for respective data distributions as indicated in figure legends. P values < 0.05 were considered statistically significant. The analyses were non-blinded.

## KEY RESOURCES TABLE

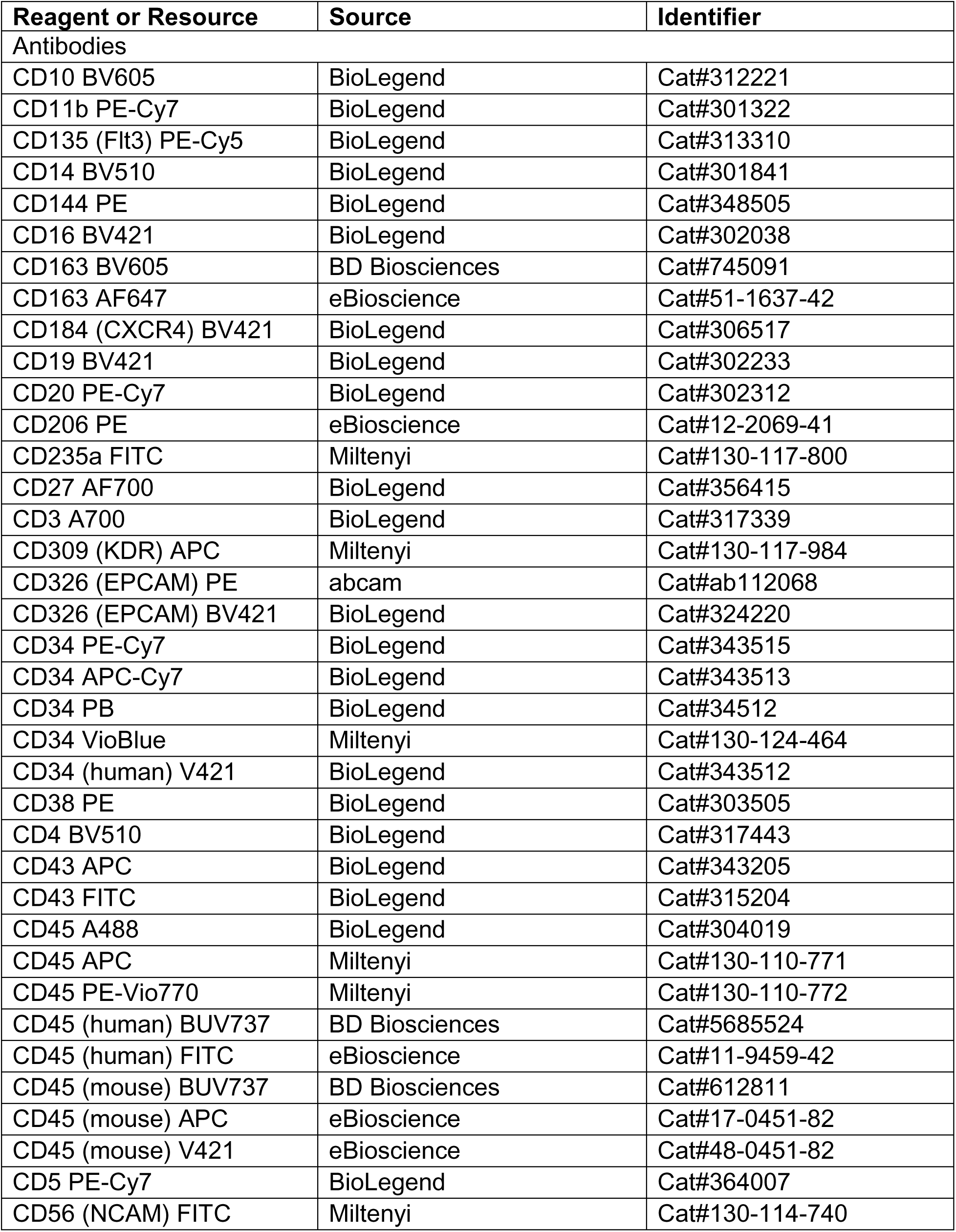
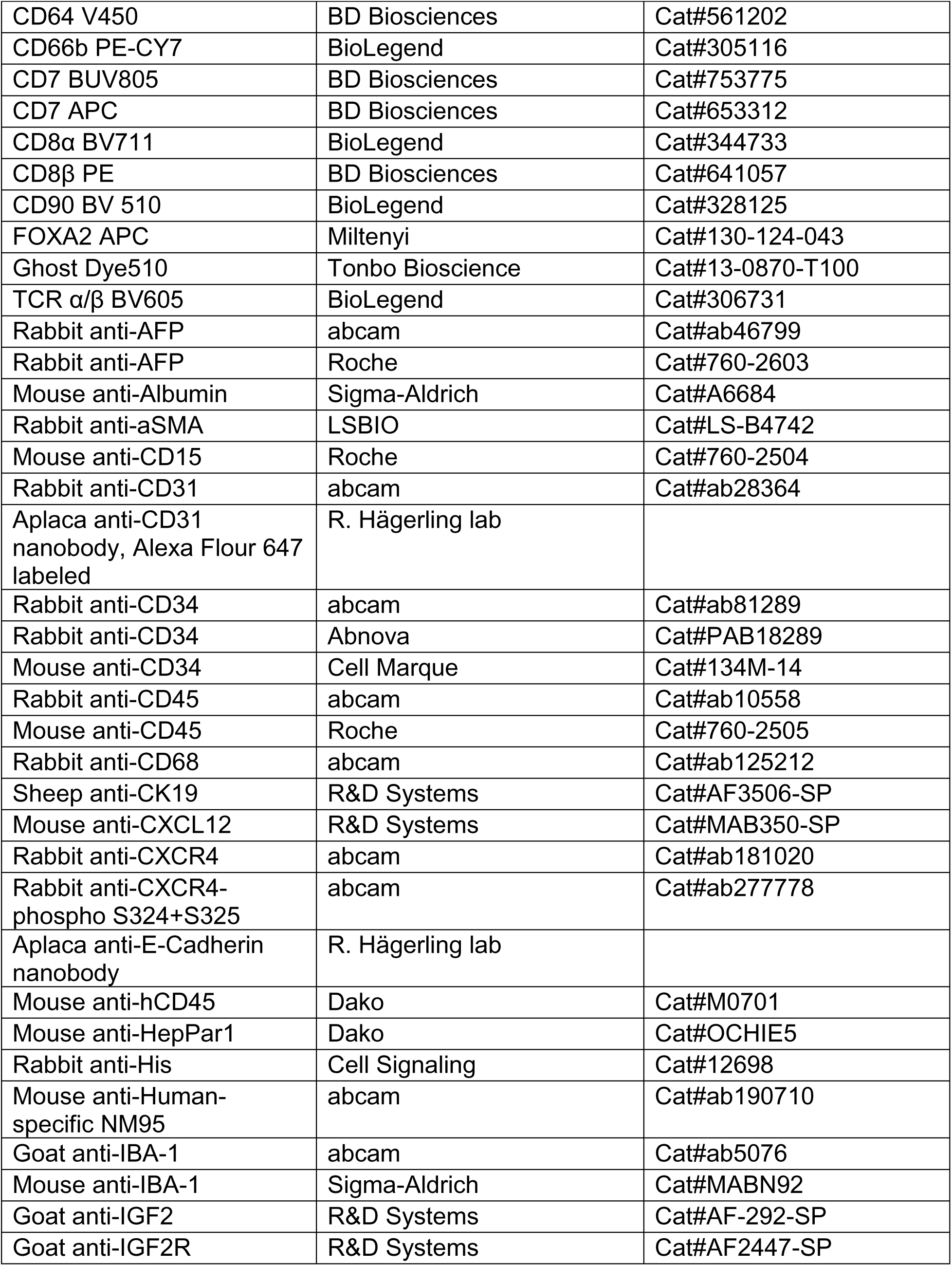
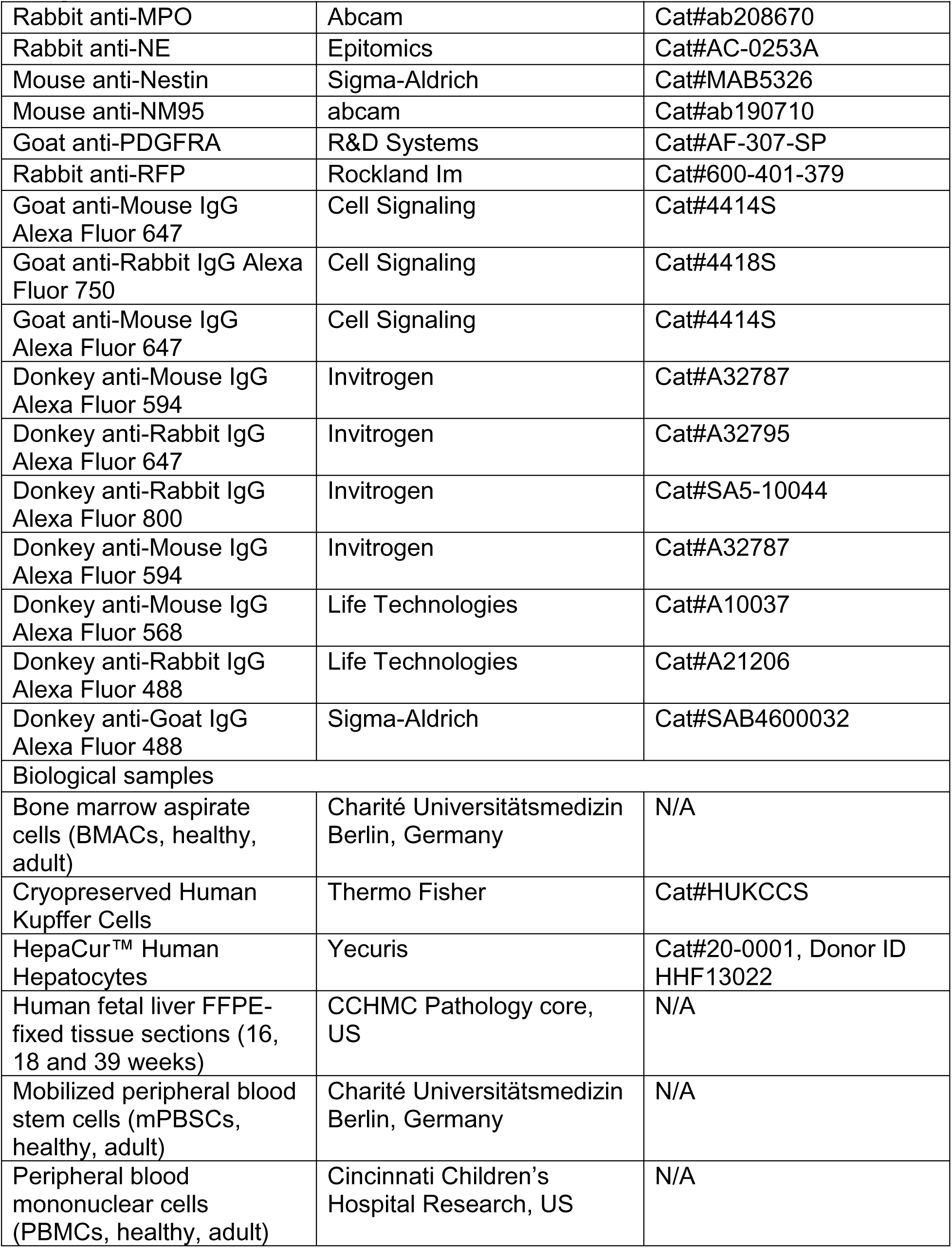
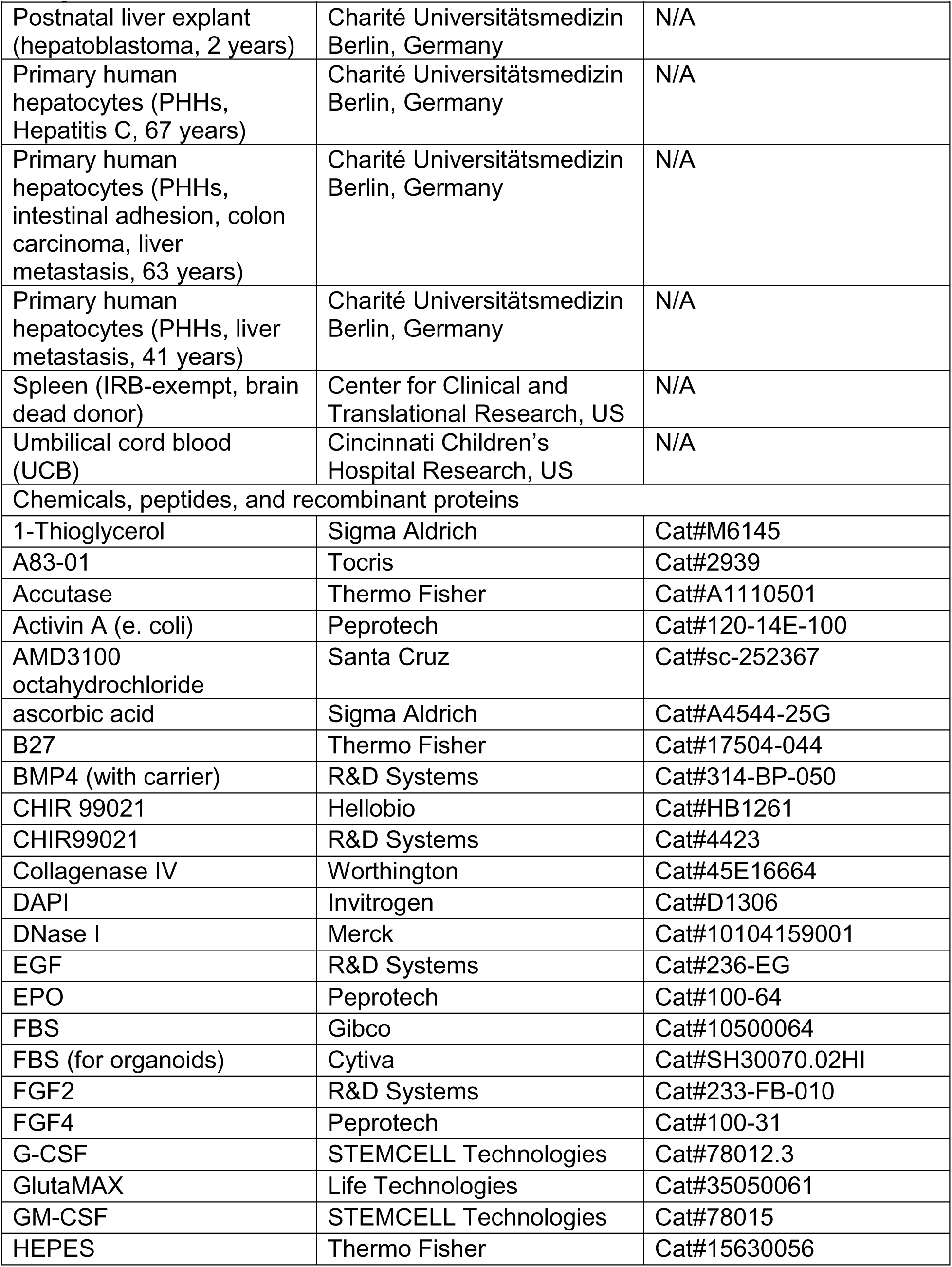
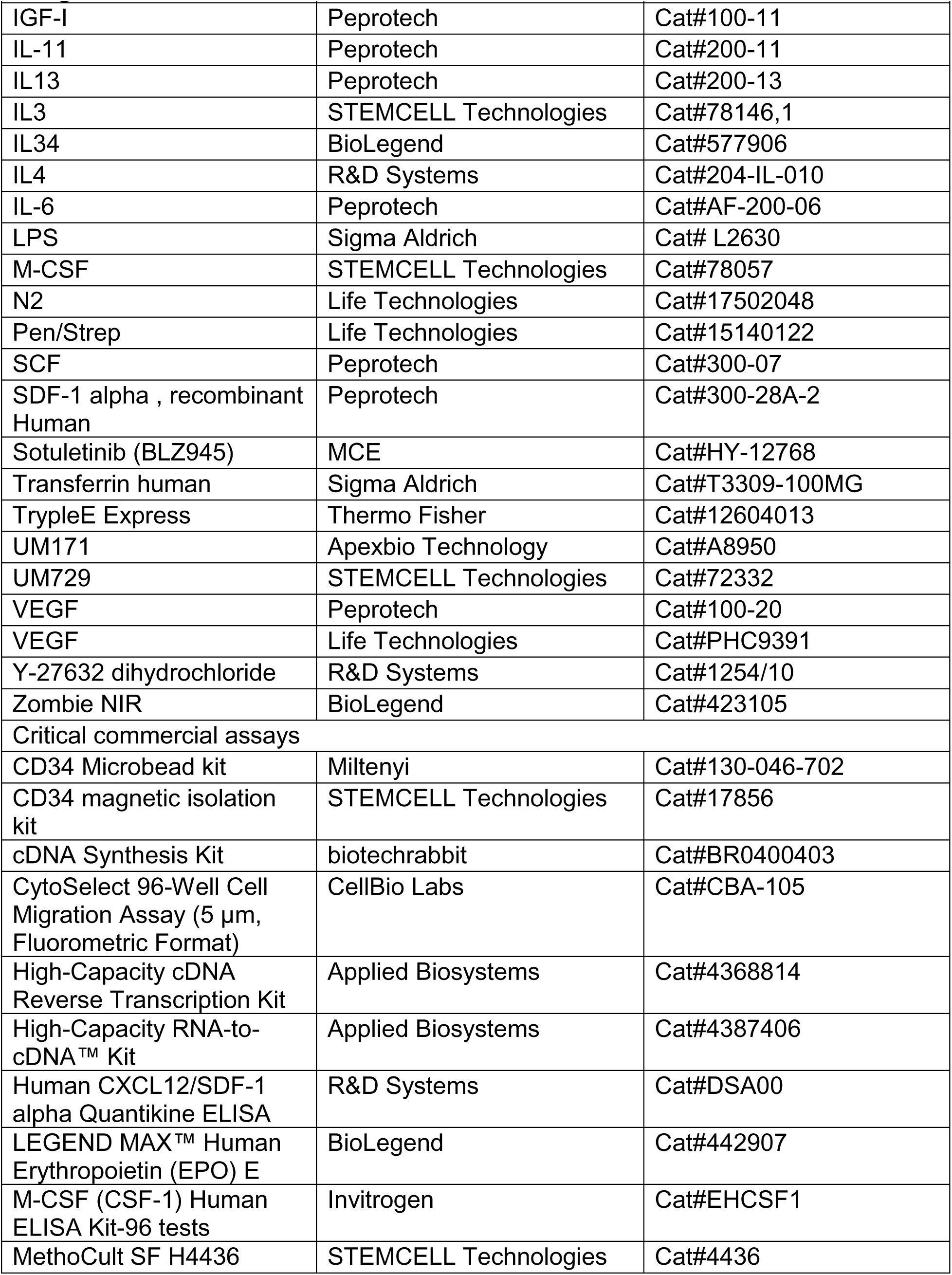
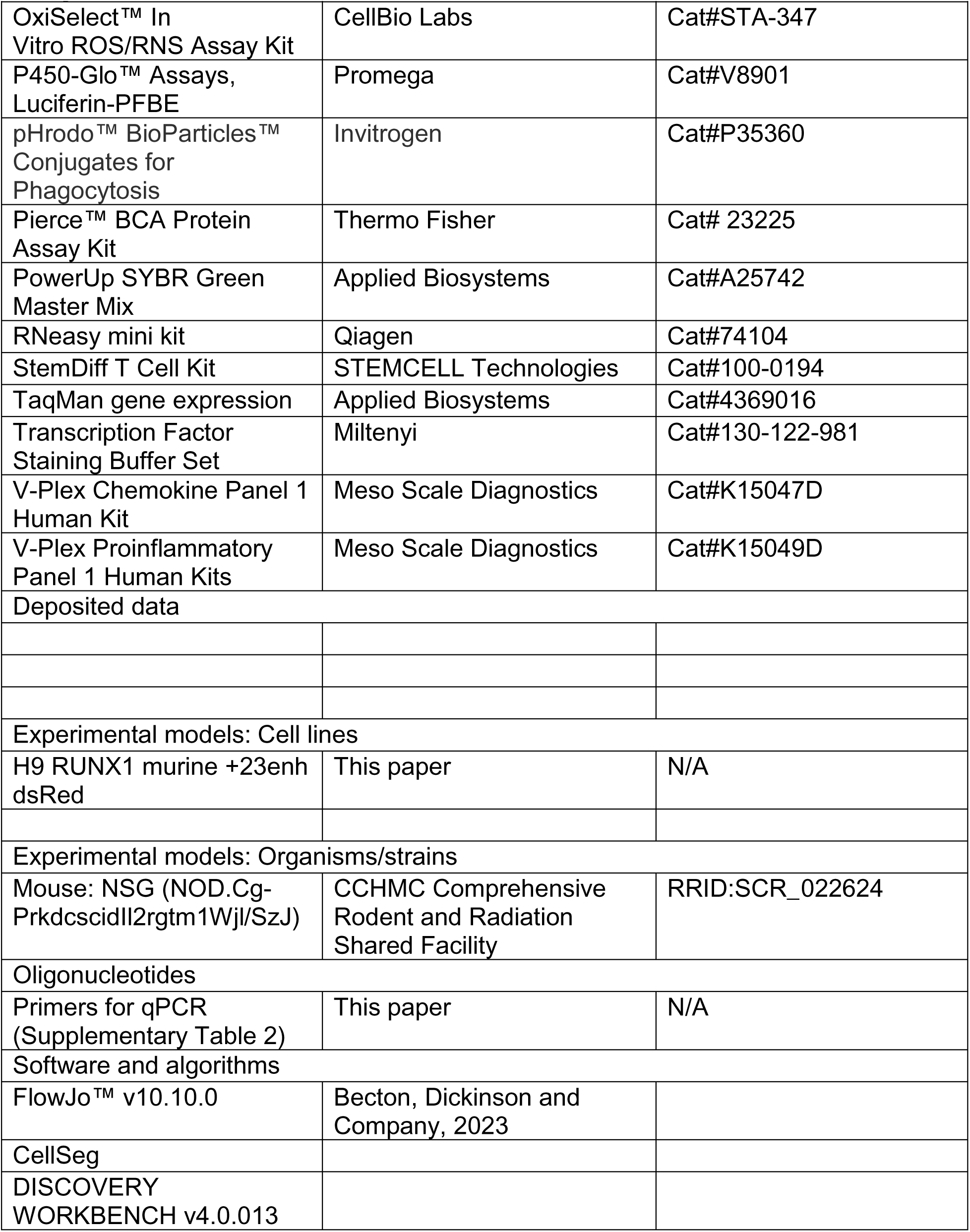
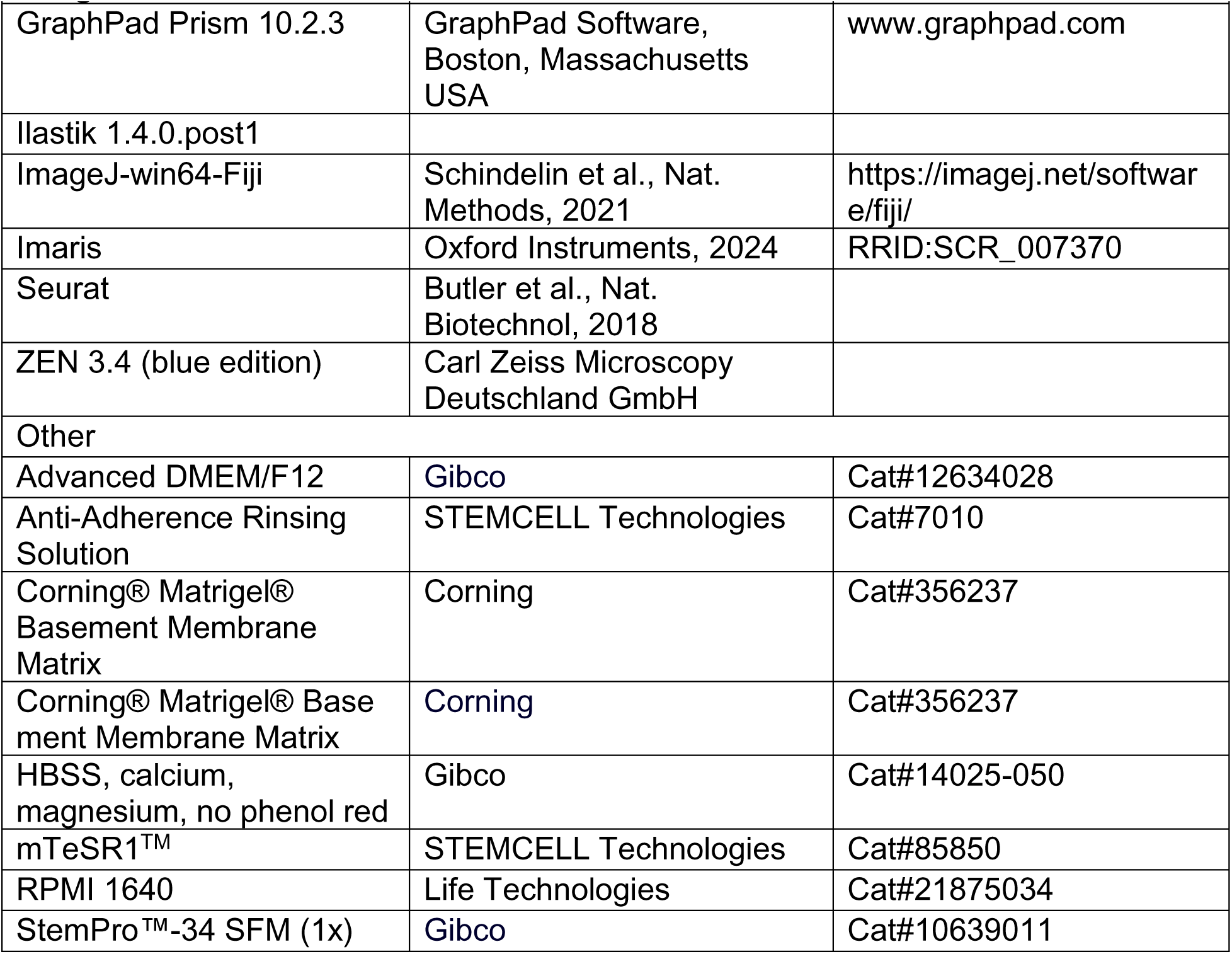

| Reagent or Resource | Source | Identifier |
| --- | --- | --- |
| Antibodies |  |  |
| CD10 BV605 | BioLegend | Cat#312221 |
| CD11b PE-Cy7 | BioLegend | Cat#301322 |
| CD135 (Flt3) PE-Cy5 | BioLegend | Cat#313310 |
| CD14 BV510 | BioLegend | Cat#301841 |
| CD144 PE | BioLegend | Cat#348505 |
| CD16 BV421 | BioLegend | Cat#302038 |
| CD163 BV605 | BD Biosciences | Cat#745091 |
| CD163 AF647 | eBioscience | Cat#51-1637-42 |
| CD184 (CXCR4) BV421 | BioLegend | Cat#306517 |
| CD19 BV421 | BioLegend | Cat#302233 |
| CD20 PE-Cy7 | BioLegend | Cat#302312 |
| CD206 PE | eBioscience | Cat#12-2069-41 |
| CD235a FITC | Miltenyi | Cat#130-117-800 |
| CD27 AF700 | BioLegend | Cat#356415 |
| CD3 A700 | BioLegend | Cat#317339 |
| CD309 (KDR) APC | Miltenyi | Cat#130-117-984 |
| CD326 (EPCAM) PE | abcam | Cat#ab112068 |
| CD326 (EPCAM) BV421 | BioLegend | Cat#324220 |
| CD34 PE-Cy7 | BioLegend | Cat#343515 |
| CD34 APC-Cy7 | BioLegend | Cat#343513 |
| CD34 PB | BioLegend | Cat#34512 |
| CD34 VioBlue | Miltenyi | Cat#130-124-464 |
| CD34 (human) V421 | BioLegend | Cat#343512 |
| CD38 PE | BioLegend | Cat#303505 |
| CD4 BV510 | BioLegend | Cat#317443 |
| CD43 APC | BioLegend | Cat#343205 |
| CD43 FITC | BioLegend | Cat#315204 |
| CD45 A488 | BioLegend | Cat#304019 |
| CD45 APC | Miltenyi | Cat#130-110-771 |
| CD45 PE-Vio770 | Miltenyi | Cat#130-110-772 |
| CD45 (human) BUV737 | BD Biosciences | Cat#5685524 |
| CD45 (human) FITC | eBioscience | Cat#11-9459-42 |
| CD45 (mouse) BUV737 | BD Biosciences | Cat#612811 |
| CD45 (mouse) APC | eBioscience | Cat#17-0451-82 |
| CD45 (mouse) V421 | eBioscience | Cat#48-0451-82 |
| CD5 PE-Cy7 | BioLegend | Cat#364007 |
| CD56 (NCAM) FITC | Miltenyi | Cat#130-114-740 |
| CD64 V450 | BD Biosciences | Cat#561202 |
| CD66b PE-CY7 | BioLegend | Cat#305116 |
| CD7 BUV805 | BD Biosciences | Cat#753775 |
| CD7 APC | BD Biosciences | Cat#653312 |
| CD8α BV711 | BioLegend | Cat#344733 |
| CD8β PE | BD Biosciences | Cat#641057 |
| CD90 BV 510 | BioLegend | Cat#328125 |
| FOXA2 APC | Miltenyi | Cat#130-124-043 |
| Ghost Dye510 | Tonbo Bioscience | Cat#13-0870-T100 |
| TCR $\alpha/\beta$ BV605 | BioLegend | Cat#306731 |
| Rabbit anti-AFP | abcam | Cat#ab46799 |
| Rabbit anti-AFP | Roche | Cat#760-2603 |
| Mouse anti-Albumin | Sigma-Aldrich | Cat#A6684 |
| Rabbit anti-aSMA | LSBIO | Cat#LS-B4742 |
| Mouse anti-CD15 | Roche | Cat#760-2504 |
| Rabbit anti-CD31 | abcam | Cat#ab28364 |
| Aplaca anti-CD31 nanobody, Alexa Flour 647 labeled | R. Hägerling lab |  |
| Rabbit anti-CD34 | abcam | Cat#ab81289 |
| Rabbit anti-CD34 | Abnova | Cat#PAB18289 |
| Mouse anti-CD34 | Cell Marque | Cat#134M-14 |
| Rabbit anti-CD45 | abcam | Cat#ab10558 |
| Mouse anti-CD45 | Roche | Cat#760-2505 |
| Rabbit anti-CD68 | abcam | Cat#ab125212 |
| Sheep anti-CK19 | R&D Systems | Cat#AF3506-SP |
| Mouse anti-CXCL12 | R&D Systems | Cat#MAB350-SP |
| Rabbit anti-CXCR4 | abcam | Cat#ab181020 |
| Rabbit anti-CXCR4-phospho S324+S325 | abcam | Cat#ab277778 |
| Aplaca anti-E-Cadherin nanobody | R. Hägerling lab |  |
| Mouse anti-hCD45 | Dako | Cat#M0701 |
| Mouse anti-HepPar1 | Dako | Cat#OCHIE5 |
| Rabbit anti-His | Cell Signaling | Cat#12698 |
| Mouse anti-Human-specific NM95 | abcam | Cat#ab190710 |
| Goat anti-IBA-1 | abcam | Cat#ab5076 |
| Mouse anti-IBA-1 | Sigma-Aldrich | Cat#MABN92 |
| Goat anti-IGF2 | R&D Systems | Cat#AF-292-SP |
| Goat anti-IGF2R | R&D Systems | Cat#AF2447-SP |
| Rabbit anti-MPO | Abcam | Cat#ab208670 |
| Rabbit anti-NE | Epitomics | Cat#AC-0253A |
| Mouse anti-Nestin | Sigma-Aldrich | Cat#MAB5326 |
| Mouse anti-NM95 | abcam | Cat#ab190710 |
| Goat anti-PDGFRα | R&D Systems | Cat#AF-307-SP |
| Rabbit anti-RFP | Rockland Im | Cat#600-401-379 |
| Goat anti-Mouse IgG<br>Alexa Fluor 647 | Cell Signaling | Cat#4414S |
| Goat anti-Rabbit IgG Alexa<br>Fluor 750 | Cell Signaling | Cat#4418S |
| Goat anti-Mouse IgG<br>Alexa Fluor 647 | Cell Signaling | Cat#4414S |
| Donkey anti-Mouse IgG<br>Alexa Fluor 594 | Invitrogen | Cat#A32787 |
| Donkey anti-Rabbit IgG<br>Alexa Fluor 647 | Invitrogen | Cat#A32795 |
| Donkey anti-Rabbit IgG<br>Alexa Fluor 800 | Invitrogen | Cat#SA5-10044 |
| Donkey anti-Mouse IgG<br>Alexa Fluor 594 | Invitrogen | Cat#A32787 |
| Donkey anti-Mouse IgG<br>Alexa Fluor 568 | Life Technologies | Cat#A10037 |
| Donkey anti-Rabbit IgG<br>Alexa Fluor 488 | Life Technologies | Cat#A21206 |
| Donkey anti-Goat IgG<br>Alexa Fluor 488 | Sigma-Aldrich | Cat#SAB4600032 |
| Biological samples |  |  |
| Bone marrow aspirate<br>cells (BMACs, healthy,<br>adult) | Charité Universitätsmedizin<br>Berlin, Germany | N/A |
| Cryopreserved Human<br>Kupffer Cells | Thermo Fisher | Cat#HUKCCS |
| HepaCur™ Human<br>Hepatocytes | Yecuris | Cat#20-0001, Donor ID<br>HHF13022 |
| Human fetal liver FFPE-fixed tissue sections (16,<br>18 and 39 weeks) | CCHMC Pathology core,<br>US | N/A |
| Mobilized peripheral blood<br>stem cells (mPBSCs,<br>healthy, adult) | Charité Universitätsmedizin<br>Berlin, Germany | N/A |
| Peripheral blood<br>mononuclear cells<br>(PBMCs, healthy, adult) | Cincinnati Children's<br>Hospital Research, US | N/A |

| **Reagent or Resource** | **Source** | **Identifier** |
| --- | --- | --- |
| Postnatal liver explant (hepatoblastoma, 2 years) | Charité Universitätsmedizin Berlin, Germany | N/A |
| Primary human hepatocytes (PHHs, Hepatitis C, 67 years) | Charité Universitätsmedizin Berlin, Germany | N/A |
| Primary human hepatocytes (PHHs, intestinal adhesion, colon carcinoma, liver metastasis, 63 years) | Charité Universitätsmedizin Berlin, Germany | N/A |
| Primary human hepatocytes (PHHs, liver metastasis, 41 years) | Charité Universitätsmedizin Berlin, Germany | N/A |
| Spleen (IRB-exempt, brain dead donor) | Center for Clinical and Translational Research, US | N/A |
| Umbilical cord blood (UCB) | Cincinnati Children's Hospital Research, US | N/A |
| Chemicals, peptides, and recombinant proteins |  |  |
| 1-Thioglycerol | Sigma Aldrich | Cat#M6145 |
| A83-01 | Tocris | Cat#2939 |
| Accutase | Thermo Fisher | Cat#A1110501 |
| Activin A (e. coli) | Peprtech | Cat#120-14E-100 |
| AMD3100 octahydrochloride | Santa Cruz | Cat#sc-252367 |
| ascorbic acid | Sigma Aldrich | Cat#A4544-25G |
| B27 | Thermo Fisher | Cat#17504-044 |
| BMP4 (with carrier) | R&D Systems | Cat#314-BP-050 |
| CHIR 99021 | Hellobio | Cat#HB1261 |
| CHIR99021 | R&D Systems | Cat#4423 |
| Collagenase IV | Worthington | Cat#45E16664 |
| DAPI | Invitrogen | Cat#D1306 |
| DNase I | Merck | Cat#10104159001 |
| EGF | R&D Systems | Cat#236-EG |
| EPO | Peprtech | Cat#100-64 |
| FBS | Gibco | Cat#10500064 |
| FBS (for organoids) | Cytiva | Cat#SH30070.02HI |
| FGF2 | R&D Systems | Cat#233-FB-010 |
| FGF4 | Peprtech | Cat#100-31 |
| G-CSF | STEMCELL Technologies | Cat#78012.3 |
| GlutaMAX | Life Technologies | Cat#35050061 |
| GM-CSF | STEMCELL Technologies | Cat#78015 |
| HEPES | Thermo Fisher | Cat#15630056 |
| IGF-I | Peprotech | Cat#100-11 |
| IL-11 | Peprotech | Cat#200-11 |
| IL13 | Peprotech | Cat#200-13 |
| IL3 | STEMCELL Technologies | Cat#78146,1 |
| IL34 | BioLegend | Cat#577906 |
| IL4 | R&D Systems | Cat#204-IL-010 |
| IL-6 | Peprotech | Cat#AF-200-06 |
| LPS | Sigma Aldrich | Cat# L2630 |
| M-CSF | STEMCELL Technologies | Cat#78057 |
| N2 | Life Technologies | Cat#17502048 |
| Pen/Strep | Life Technologies | Cat#15140122 |
| SCF | Peprotech | Cat#300-07 |
| SDF-1 alpha , recombinant Human | Peprotech | Cat#300-28A-2 |
| Sotuletinib (BLZ945) | MCE | Cat#HY-12768 |
| Transferrin human | Sigma Aldrich | Cat#T3309-100MG |
| TrypleE Express | Thermo Fisher | Cat#12604013 |
| UM171 | Apexbio Technology | Cat#A8950 |
| UM729 | STEMCELL Technologies | Cat#72332 |
| VEGF | Peprotech | Cat#100-20 |
| VEGF | Life Technologies | Cat#PHC9391 |
| Y-27632 dihydrochloride | R&D Systems | Cat#1254/10 |
| Zombie NIR | BioLegend | Cat#423105 |
| Critical commercial assays |  |  |
| CD34 Microbead kit | Miltenyi | Cat#130-046-702 |
| CD34 magnetic isolation kit | STEMCELL Technologies | Cat#17856 |
| cDNA Synthesis Kit | biotechrabbit | Cat#BR0400403 |
| CytoSelect 96-Well Cell Migration Assay (5 µm, Fluorometric Format) | CellBio Labs | Cat#CBA-105 |
| High-Capacity cDNA Reverse Transcription Kit | Applied Biosystems | Cat#4368814 |
| High-Capacity RNA-to-cDNA™ Kit | Applied Biosystems | Cat#4387406 |
| Human CXCL12/SDF-1 alpha Quantikine ELISA | R&D Systems | Cat#DSA00 |
| LEGEND MAX™ Human Erythropoietin (EPO) E | BioLegend | Cat#442907 |
| M-CSF (CSF-1) Human ELISA Kit-96 tests | Invitrogen | Cat#EHCSF1 |
| MethoCult SF H4436 | STEMCELL Technologies | Cat#4436 |
| OxiSelect™ In Vitro ROS/RNS Assay Kit | CellBio Labs | Cat#STA-347 |
| P450-Glo™ Assays, Luciferin-PFBE | Promega | Cat#V8901 |
| pHrodo™ BioParticles™ Conjugates for Phagocytosis | Invitrogen | Cat#P35360 |
| Pierce™ BCA Protein Assay Kit | Thermo Fisher | Cat# 23225 |
| PowerUp SYBR Green Master Mix | Applied Biosystems | Cat#A25742 |
| RNeasy mini kit | Qiagen | Cat#74104 |
| StemDiff T Cell Kit | STEMCELL Technologies | Cat#100-0194 |
| TaqMan gene expression | Applied Biosystems | Cat#4369016 |
| Transcription Factor Staining Buffer Set | Miltenyi | Cat#130-122-981 |
| V-Plex Chemokine Panel 1 Human Kit | Meso Scale Diagnostics | Cat#K15047D |
| V-Plex Proinflammatory Panel 1 Human Kits | Meso Scale Diagnostics | Cat#K15049D |
| Deposited data |  |  |
| Experimental models: Cell lines |  |  |
| H9 RUNX1 murine +23enh dsRed | This paper | N/A |
| Experimental models: Organisms/strains |  |  |
| Mouse: NSG (NOD.Cg-Prkdcscidll2rgtm1Wjl/SzJ) | CCHMC Comprehensive Rodent and Radiation Shared Facility | RRID:SCR_022624 |
| Oligonucleotides |  |  |
| Primers for qPCR (Supplementary Table 2) | This paper | N/A |
| Software and algorithms |  |  |
| FlowJo™ v10.10.0 | Becton, Dickinson and Company, 2023 |  |
| CellSeg |  |  |
| DISCOVERY WORKBENCH v4.0.013 |  |  |
| GraphPad Prism 10.2.3 | GraphPad Software, Boston, Massachusetts USA | www.graphpad.com |
| Ilastik 1.4.0.post1 |  |  |
| ImageJ-win64-Fiji | Schindelin et al., Nat. Methods, 2021 | https://imagej.net/software/fiji/ |
| Imaris | Oxford Instruments, 2024 | RRID:SCR_007370 |
| Seurat | Butler et al., Nat. Biotechnol, 2018 |  |
| ZEN 3.4 (blue edition) | Carl Zeiss Microscopy Deutschland GmbH |  |
| Other |  |  |
| Advanced DMEM/F12 | Gibco | Cat#12634028 |
| Anti-Adherence Rinsing Solution | STEMCELL Technologies | Cat#7010 |
| Corning® Matrigel® Basement Membrane Matrix | Corning | Cat#356237 |
| Corning® Matrigel® Basement Membrane Matrix | Corning | Cat#356237 |
| HBSS, calcium, magnesium, no phenol red | Gibco | Cat#14025-050 |
| mTeSR1™ | STEMCELL Technologies | Cat#85850 |
| RPMI 1640 | Life Technologies | Cat#21875034 |
| StemPro™-34 SFM (1x) | Gibco | Cat#10639011 |

## ACKNOWLEDGEMENTS

The authors of this scientific article would also like to acknowledge the funding provided by the Deutsche Forschungsgemeinschaft (Emmy Noether Programme) and Eva Luise und Horst Köhler Stiftung (Alliance4Rare Program) to M.R. This work was also supported by Cincinnati Children’s Research Foundation grant, the Falk Transformational Awards Program, NIH Director’s New Innovator Award (DP2 DK128799-01) and CREST (20gm1210012h0001) grant from Japan Agency for Medical Research and Development (AMED) to TT. This work was also supported by an NIH grant UG3/UH3 DK119982, Cincinnati Center for Autoimmune Liver Disease Fellowship Award, PHS Grant P30 DK078392 (Integrative Morphology Core and Pluripotent Stem Cell and Organoid Core) of the Digestive Disease Research Core Center in Cincinnati, Takeda Science Foundation Award, Mitsubishi Foundation Award and AMED grants JP18fk0210037h0001, JP18bm0704025h0001, JP21gm1210012h0002, JP21bm0404045h0003, and JP21fk0210060h0003, JST Moonshot JPMJMS2022-10 and JPMJMS2033-12, and JSPS KAKENHI Grant JP18H02800, 19K22416. TT is a New York Stem Cell Foundation – Robertson Investigator. KI is a New York Stem Cell Foundation – Druckenmiller fellow. The authors of this scientific article would also like to acknowledge the funding provided by the ERC starting grant 2019 ‘EpiTune’ (803992) and DFG project (PO 2058/3-1) to JKP. We would like to extend our gratitude to David Grier of CCHMC for his contributions to the hemopathological analysis. Additionally, we would like to thank Harald Stachelscheid and Katarzyna Ludwik of the BIH Core Unit pluripotent Stem cells and Organoids (CUSCO) for sharing expertise with iPSC-culture and differentiation of iHeps. We thank the BIH Flow and Mass Cytometry Core, the CCHMC Pluripotent Stem Cell Facility (PSCF, RRID: SCR_022634), Bio-Imaging and Analysis Facility (BAF, RRID: SCR_022628), Transgenic Animal, Genome Editing Facility (TAGE, RRID: SCR_022642), Comprehensive Rodent and Radiation Facility (CRRF, RRID: SCR_022624) and Single Cell Genomics Facility (SCGF, RRID: SCR_022653). We are also grateful for the advice provided by Mark Wunderlich (Mulloy Lab, CCHMC) and the support provided by the members of the Takebe Lab, Zorn Lab, Wells Lab, Cancelas Lab at CCHMC. We would like to thank the CCHMC Research Flow Cytometry Facility (RFCF, RRID: SCR_022635, supported by NIH S10OD023410), and the Integrated Pathology Research Facility (IPRF, RRID:SCR_022637) for their contributions, including providing FACS experiments and support with immunostainings and providing fetal specimens, respectively. We thank Prof. Dr. med. Frank Tacke for his expertise and his laboratory for access to resources. We thank Prof. Johannes Schulte at Charité Berlin for sharing human CD34+ and bone marrow aspirate-derived cells from his laboratory, Prof. James Mulloy at CCHMC for sharing human CD34+ cells from his laboratory, and Prof. Dr. med. Igor Sauer and laboratory (Charité Berlin) for sharing primary human hepatocytes – all after obtaining informed consent. We thank Prof. Dr. rer. nat. Simon Haas, Dr. rer. nat. Marlene-Sophia Kohlhepp and Guo Yin for sharing their expertise. We thank former Rezvani Lab members Natalia Martagon-Calderon, Ferhat Ali Yaman, Pia Dernick, and former Bufler Lab (Charité Berlin) member Kerstin Sommer for technical assistance. We extend our deepest gratitude to the courageous families who generously consented to provide blood-derived cells or donated their loved one’s organs and tissues for biomedical research (*i.e.* spleen). Such important research like this would not be possible without this selfless gift of hope. Thanks to Dr. Appakalai N. Balamurugan’s team and the organ procurement organizations, Kentucky Organ Donor Affiliates (KODA), Louisville; LifeCenter, Cincinnati; and Lifeline of Ohio, Columbus for supporting these special families.

## ABBREVIATIONS

AGM: Aorta–Gonad–Mesonephros
AH: Adult Hepatocyte
BMAC: Bone Marrow Aspirate Cell
CFU: Colony Forming Units
CS: Carnegie Stage
EHT: Endothelial-to-Hematopoietic-Transition
ESC: Embryonic Stem Cell
FL: Fetal Liver
FLO: Human Fetal Liver-like Organoid
GMP: Granulocyte–Monocyte Progenitor
HB: Hemoglobin
HPC: Hematopoietic Progenitors Cell
HSC: Hematopoietic Stem Cell
HSPC: Hematopoietic Stem and Progenitor Cell
IHC: Immunohistochemistry
iPSC: induced Pluripotent Stem Cell
mPBSC: mobilized Peripheral Blood Stem Cell
mIF: Multiplex Immunofluorescence
pcw: Post conception week
PBMC: Peripheral Blood Mononuclear Cell
PSC: Pluripotent Stem Cell
scRNA-Seq: Single-cell RNA sequencing
UCB: Umbilical Cord Blood
YS: Yolk Sac

## Notes

### Competing Interest Statement

The authors have declared no competing interest.

