## Supplementary figures and images for "Fetal Liver-like Organoids Recapitulate Blood-Liver Niche Development and Multipotent Hematopoiesis from Human Pluripotent Stem Cells"

### Supplementary Figures

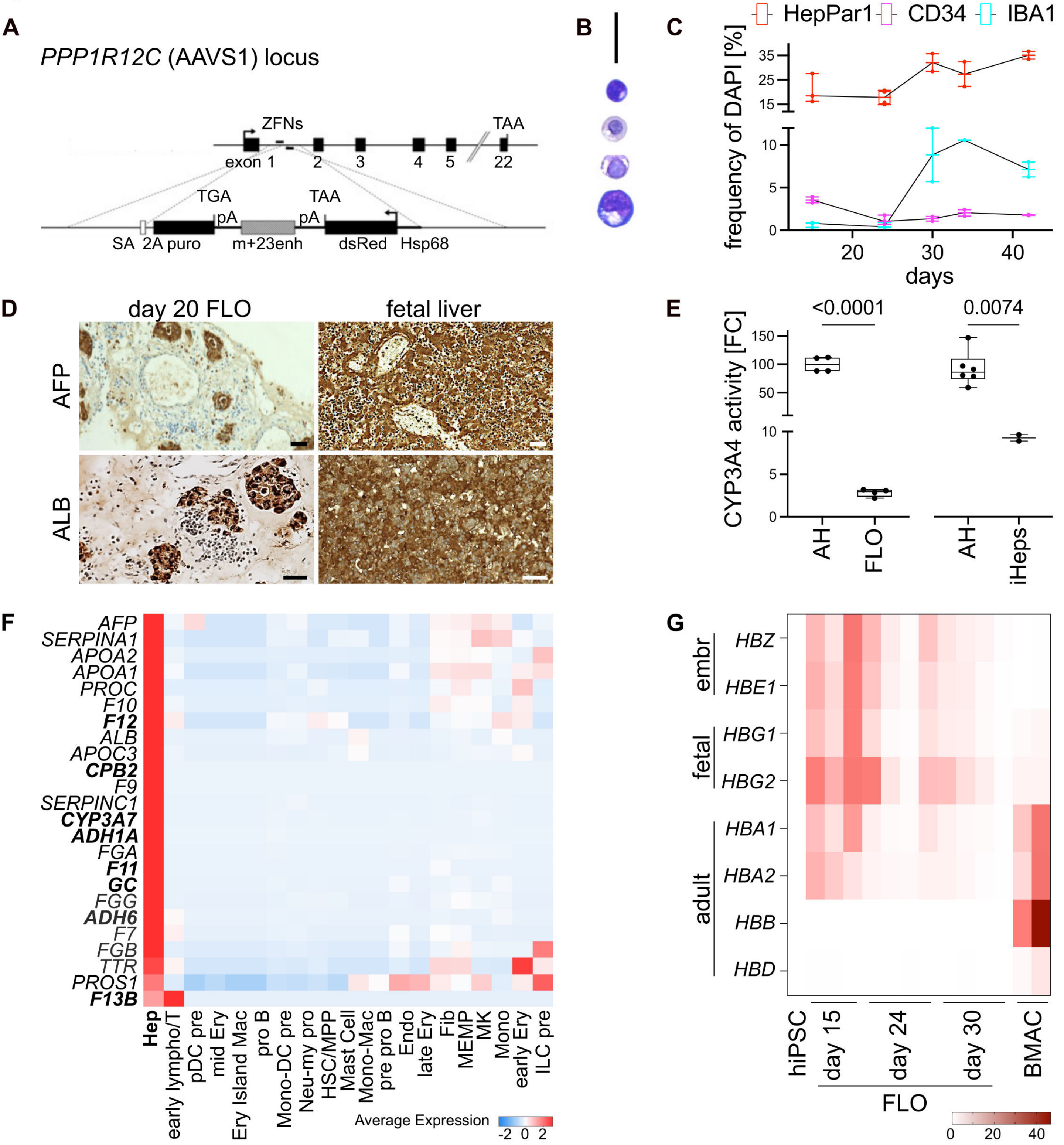

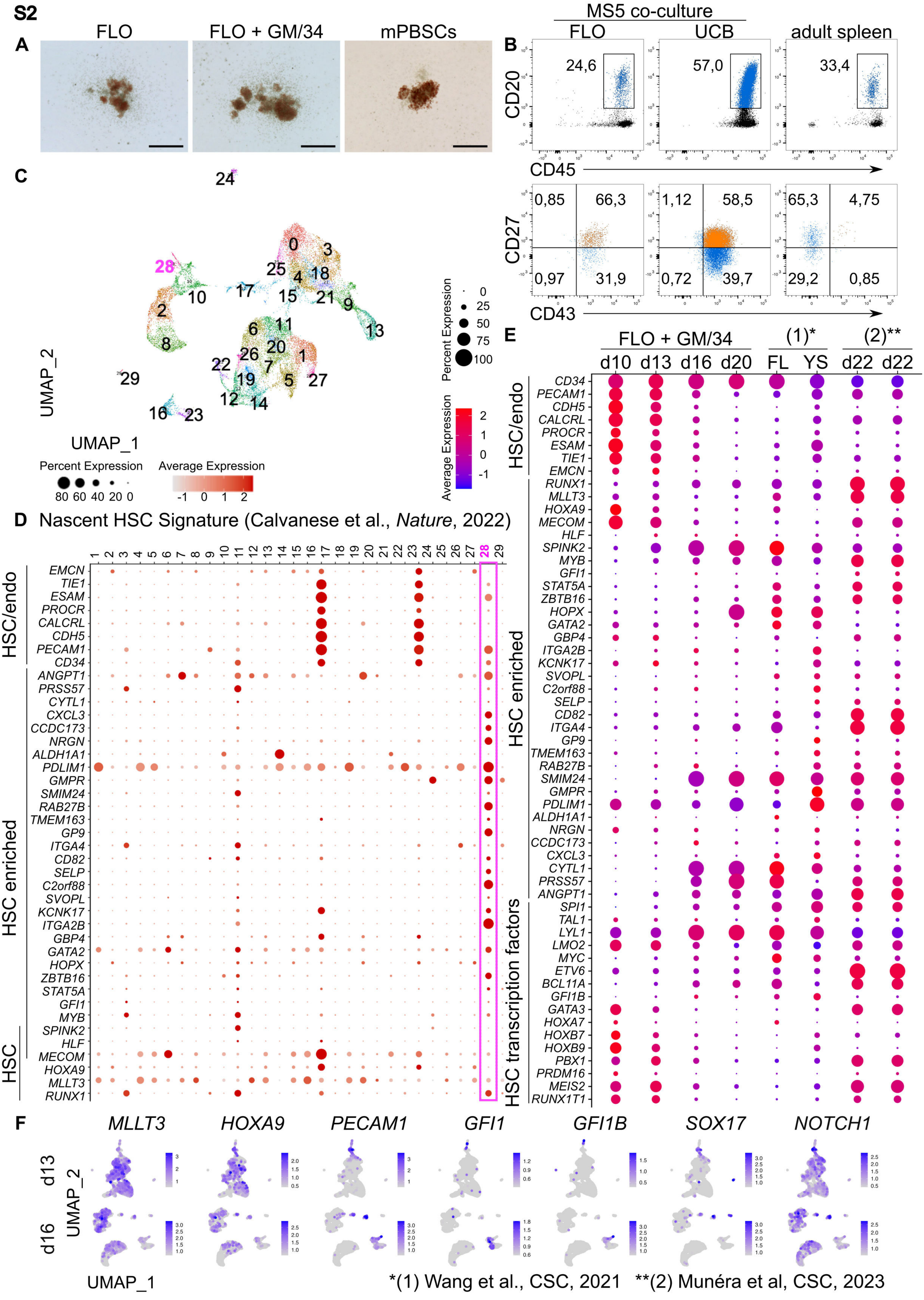

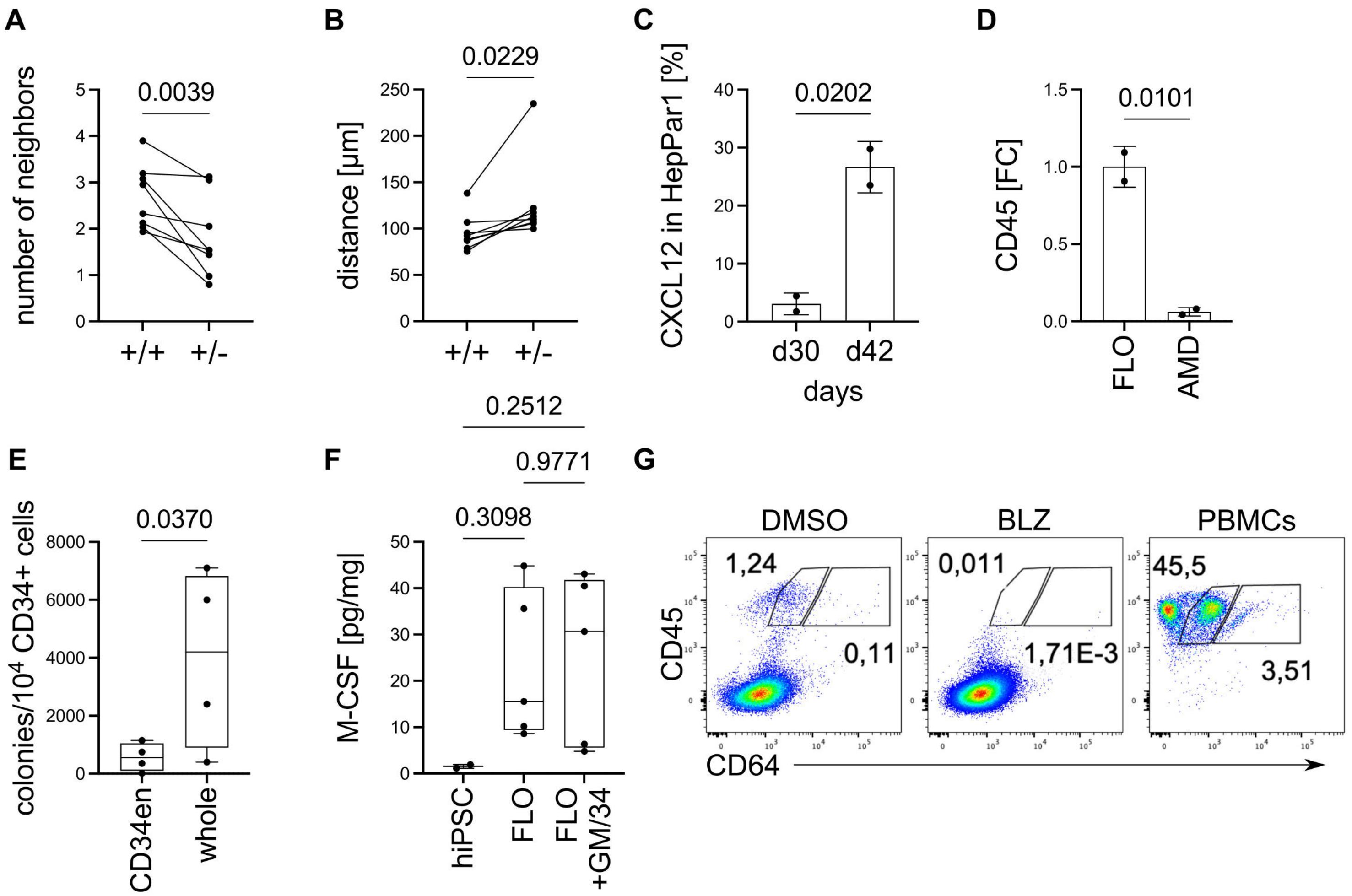

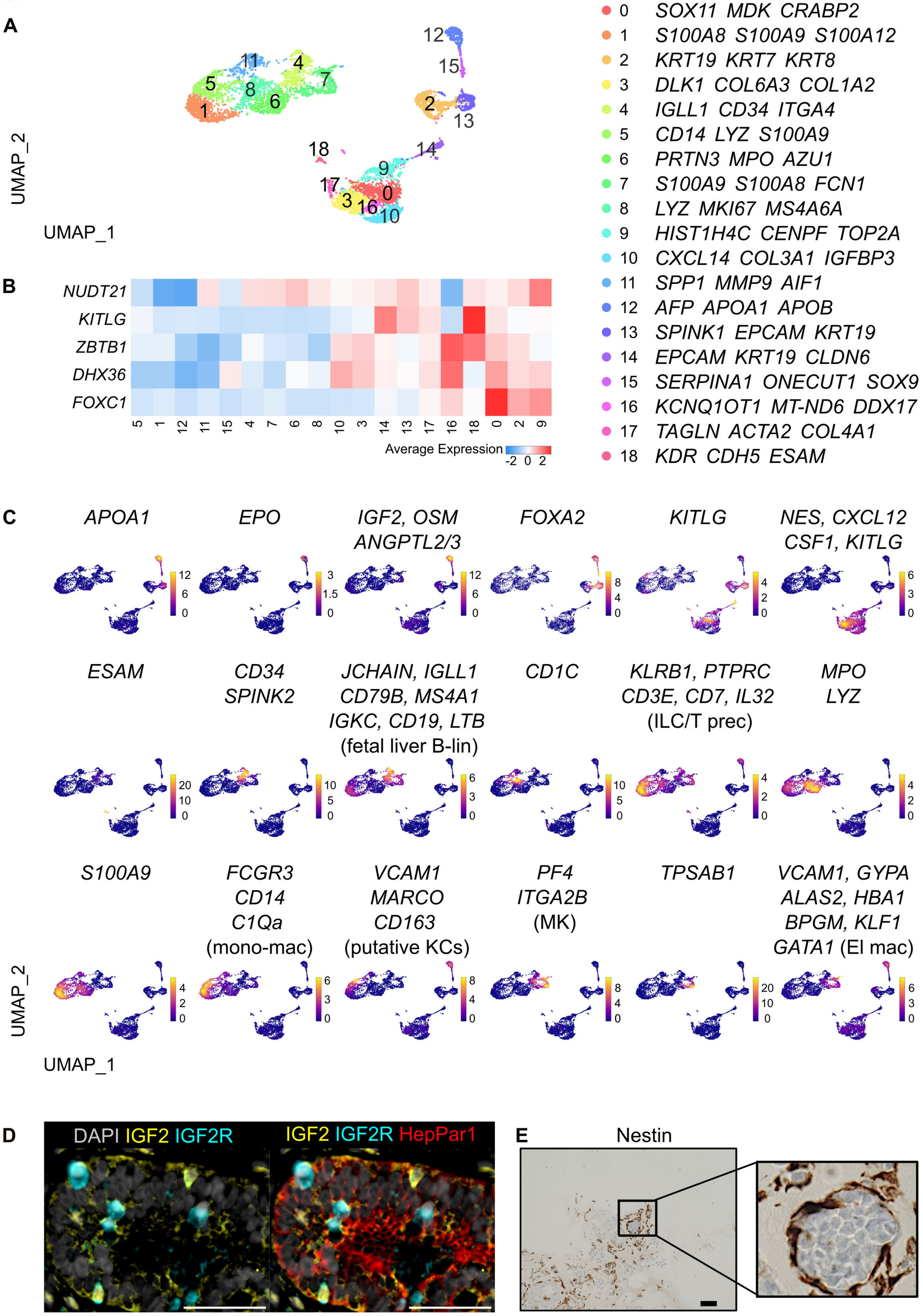

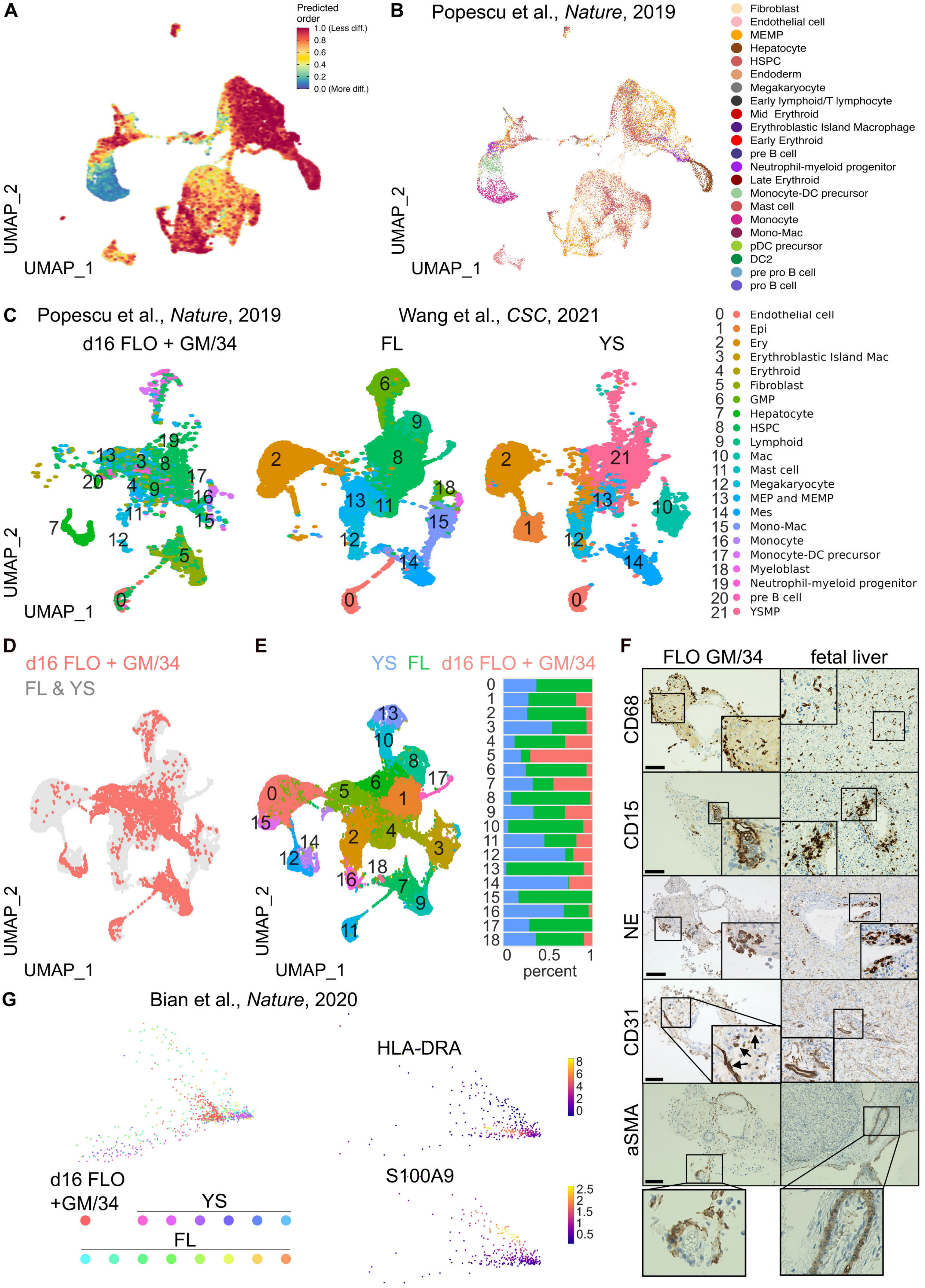

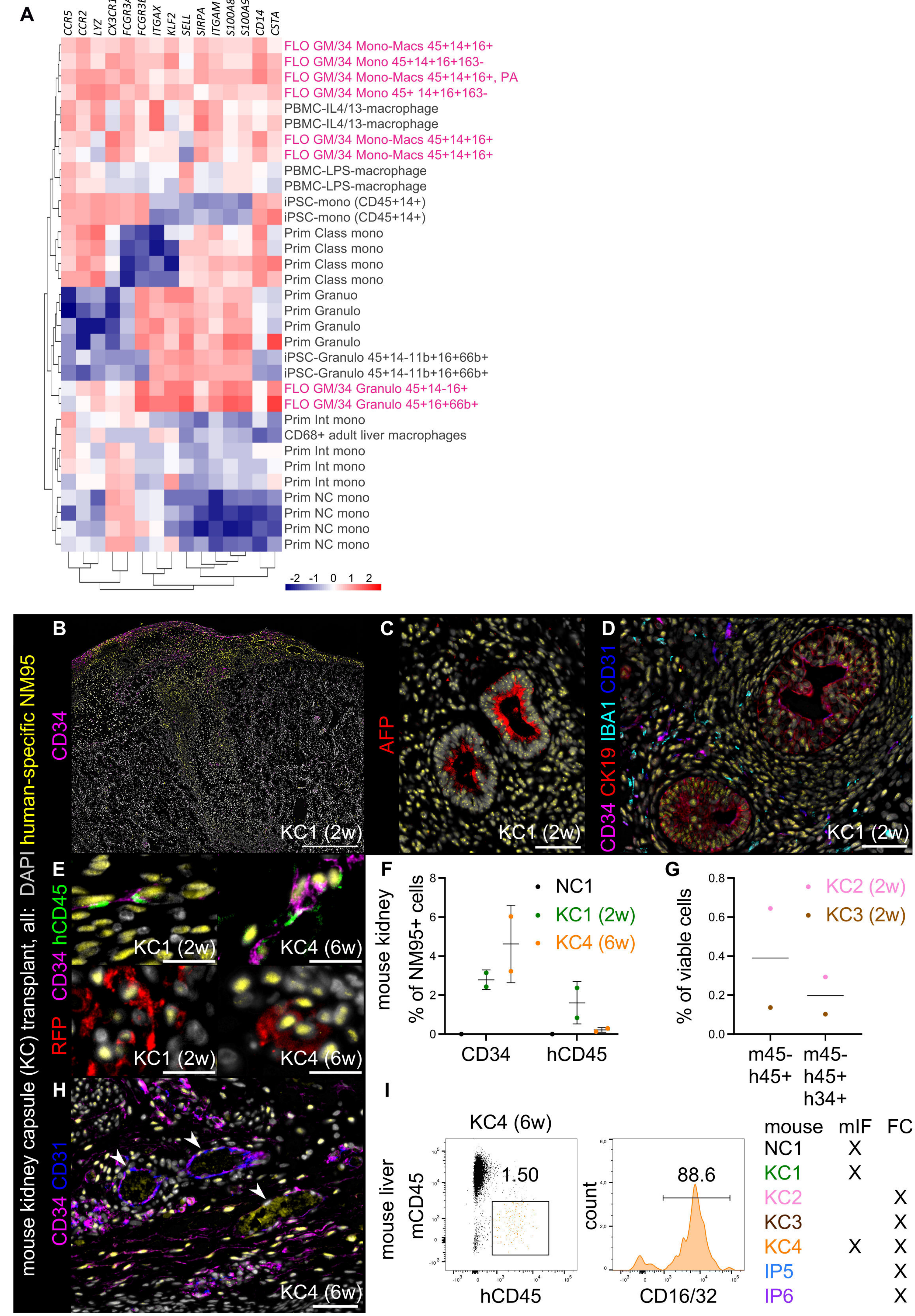
